# A phosphorylation signal activates genome-wide transcriptional control by BfmR, the global regulator of *Acinetobacter* resistance and virulence

**DOI:** 10.1101/2024.06.16.599214

**Authors:** Nicole Raustad, Yunfei Dai, Akira Iinishi, Arpita Mohapatra, Mark W. Soo, Everett Hay, Gabrielle M. Hernandez, Edward Geisinger

## Abstract

The nosocomial pathogen *Acinetobacter baumannii* is a major threat to human health. The sensor kinase-response regulator system, BfmS-BfmR, is essential to multidrug resistance and virulence in the bacterium and represents a potential antimicrobial target. Important questions remain about how the system controls resistance and pathogenesis. Although BfmR knockout alters expression of >1000 genes, its direct regulon is undefined. Moreover, how phosphorylation controls the regulator is unclear. Here, we address these problems by combining mutagenesis, ChIP-seq, and in vitro phosphorylation to study the functions of phospho-BfmR. We show that phosphorylation is required for BfmR-mediated gene regulation, antibiotic resistance, and sepsis development in vivo. Consistent with activating the protein, phosphorylation induces dimerization and target DNA affinity. Integrated analysis of genome-wide binding and transcriptional profiles of BfmR led to additional key findings: (1) Phosphorylation dramatically expands the number of genomic sites BfmR binds; (2) DNA recognition involves a direct repeat motif widespread across promoters; (3) BfmR directly regulates 303 genes as activator (eg, capsule, peptidoglycan, and outer membrane biogenesis) or repressor (pilus biogenesis); (4) BfmR controls several non-coding sRNAs. These studies reveal the centrality of a phosphorylation signal in driving *A. baumannii* disease and disentangle the extensive pathogenic gene-regulatory network under its control.

## INTRODUCTION

The Gram-negative pathogen *Acinetobacter baumannii* causes drug-resistant nosocomial diseases such as ventilator-associated pneumonia, wound infections, and sepsis (1). Prevalence of resistant *Acinetobacter* infections is particularly high in hospital critical care units and has increased during the COVID-19 pandemic (2,3). Given the frequency of multidrug resistance and the limited number of treatment options currently available, the problem of *A. baumannii* has been elevated to the highest level of urgency by the CDC (4). To aid the development of novel therapeutic and preventative strategies against *A. baumannii*, it is important that we understand the mechanisms allowing the pathogen to establish nosocomial infections and resist treatments.

Two-component systems (TCSs), consisting of an inner membrane sensor kinase and a cytoplasmic response regulator, are commonly used by pathogenic bacteria to respond to environmental stimuli and contribute to disease (5). In *A. baumannii*, the TCS BfmRS has an especially crucial role in controlling many aspects of pathogenicity and intrinsic antibiotic resistance (6,7). BfmRS was first identified as a regulator of biofilm formation (8). The system has since been shown to regulate a number of additional features relevant to infection, including capsule production (9,10), serum resistance (6,7), fitness and virulence in animal models (6,11,12), antibiotic resistance and tolerance (6,7,13), desiccation tolerance (14,15), and motility (16). These observations support the idea that BfmRS is essential to what makes *A. baumannii* a nosocomial pathogen and presents a potential antimicrobial target (7).

For many of the above phenotypes, knockouts of BfmR (response regulator) and BfmS (membrane sensor kinase) have opposing effects, consistent with control of BfmR activity by BfmS. For instance, deletions of *bfmR* (eg, Δ*bfmR* or Δ*bfmRS*) greatly increase susceptibility to the complement system and antibiotic treatments, whereas Δ*bfmS* lowers susceptibility to many of the same challenges (6). Opposing effects are also observed on capsule levels and transcriptional reporters targeted by BfmRS (6,17). Δ*bfmS* increases the level of phosphorylated BfmR, an effect highly similar to mutations in other loci, such as the *dsbA* envelope disulfide enzyme, that activate the system (9,17). BfmS thus appears to regulate BfmR activity by modulating its phosphorylation, and its deletion provides a genetic background that serves to mimic stress conditions activating the TCS.

In accord with the pleiotropic phenotypes BfmRS controls, RNA-seq has revealed that the system governs an expansive regulon of >1000 genes (6,17). The genes map to diverse pathways, including cell envelope synthesis (e.g., production of capsule, cell wall, and outer membrane), β-lactamases, pilus formation, and protection against oxidative and osmotic stress. A comparable number of genes within the regulon show decreased or increased expression when comparing *bfmR* knockouts to WT (6,17). These results suggest that BfmR may function as a transcriptional activator or repressor of different targets, a possibility that we examine in this paper. BfmRS is known to mediate responses to stress from cell-wall and ribosome-targeting antibiotics (10) as well as envelope disulfide formation (9), but the precise contribution of the regulon in conferring protection against these stresses is poorly understood.

Structural and biochemical analyses of BfmR have begun to shed light into its regulation mechanisms. BfmR belongs to the large OmpR/PhoB family of response regulators (7). Members of this family contain an N-terminal receiver domain and C-terminal DNA binding domain (18,19) and function as transcriptional activators, repressors, or both, depending on the homolog studied (20,21). Phosphorylation of the receiver typically causes protein homodimerization and increases target DNA affinity (20), although one or both activities can be seen in the absence of phosphorylation in some cases (22). X-ray crystallography and solution NMR of isolated BfmR domains revealed structural features shared with OmpR/PhoB proteins. These include, in the receiver domain, a similarly positioned aspartate (Asp58) predicted to serve as the site of phosphorylation and other active site residues; and in the C-terminal domain, the characteristic winged helix-turn-helix fold (23). Phosphorylation at Asp58 has not been demonstrated, although mutation and treatment with a phosphomimetic (BeF_3_^-^) have been used to examine the function of this site. A D58A mutation predicted to block phosphorylation abolished BfmR-dependent control of transcriptional reporters (17) and motility (16). BeF_3_^-^, which binds to Asp58 in the crystal structure, caused BfmR to shift from monomeric to dimeric in solution as assessed by size-exclusion chromatography (SEC) (23). BfmR interacts with DNA sequences upstream of *bfmRS* (23) and 4 additional genes (17), and BeF_3_^-^ treatment lowered affinity for the *bfmRS* promoter sequence (23). How BfmR interacts with additional DNA targets and how phosphorylation affects its activity is incompletely understood. Moreover, the full spectrum of directly regulated BfmR targets is yet to be defined.

In this paper we have addressed the above questions by profiling the interactions of BfmR with sites throughout the *A. baumannii* genome, using strains and conditions allowing control of BfmR phosphorylation both *in vivo* and *in vitro*. Our findings support Asp58 as the site of phosphorylation, confirm phosphorylation-dependent dimerization, and show that the phosphorylation signal is critical for intrinsic antibiotic resistance and virulence. Moreover, we demonstrate that BfmR binds hundreds of sites dispersed throughout the genome in a phosphorylation-dependent manner, and we use this information to define its global direct regulon. This work reveals the importance of BfmR phosphorylation in the global control of *A. baumannii* pathogenicity.

## MATERIALS AND METHODS

### Bacterial strains, growth conditions, and antibiotics

Bacterial strains used in this work are described in Table S1. *A. baumannii* strains were derivatives of ATCC 17978. All cultures were in lysogeny broth (LB) (10 g/L tryptone, 5 g/L yeast extract, 10 g/L NaCl) medium. Medium was supplemented with antibiotics as needed for isolation of strains with plasmids: Tetracycline (Tc, 10µg/mL) and Gentamicin (Gm, 10µg/mL) for *A. baumannii*; Tc (10µg/mL), Gm (30µg/mL), Kanamycin (Km, 30-50µg/ml), Carbenicillin (50-100µg/ml), Chloramphenicol (Cm, 25µg/mL) for *E. coli*. Liquid cultures were incubated at 37°C (unless otherwise indicated) with shaking in flasks or in tubes on a roller drum.

### Molecular cloning and strain construction

Plasmids and oligos used in this study are listed in Table S1. Oligos were purchased from IDT. Cloned PCR products were verified by sequencing (Genewiz). Plasmids were introduced into *A. baumannii* by electroporation. Genotypes of recombinant mutant *A. baumannii* strains isolated by allelic exchange were verified by colony PCR (all strains) and/or genome resequencing (NRA10, JBA202, NRA407, and NRA486; SeqCenter).

To construct the *A. baumannii bfmR*(D58A) strain, ∼1kb centered on codon 58 of *bfmR* from NRA29 was amplified by PCR and cloned in pUC18, resulting in pNRE157. The construct was subcloned in pJB4648 using BamHI and SalI, resulting in pNRE169. *bfmR*(D58A) recombinants were isolated in *A. baumannii* 17978 by two-step allelic exchange with sucrose counterselection as described, creating NRA407. To construct a *bfmR*(D58A) Δ*bfmS* strain, pNRE169 was introduced into EGA195 and the double recombinant (NRA446) was isolated by allele exchange as above. To construct *bfmR*(D58A) Δ*gtr6*, pNRE169 was recombined in JBA202, followed by allelic exchange to isolate NRA486.

To construct *A. baumannii* strains encoding BfmR^3XFLAG^, the BamHI-SalI fragment of pEGE228 containing *bfmR*3XFLAG-*bfmS* was subcloned in pJB4846, creating pNRE26. *bfmR*3XFLAG recombinants were isolated by allele exchange in *A. baumannii* 17978 as above, resulting in NRA28. A *bfmR*(D58A)3XFLAG-*bfmS* construct was made by inverse PCR using primers bfmR-D58A-F and bfmR-D58x-R and pEGE228 as template. The resulting product was phosphorylated and circularized by ligation, creating pNRE12. The *bfmR*(D58A)3XFLAG-*bfmS* fragment was then subcloned in pJB4648 using BamHI and SalI restriction sites, creating pNRE27. The *bfmR*(D58A)3XFLAG recombinant (NRA29) was isolated via allele exchange in NRA28. A *bfmR*3XFLAG-Δ*bfmS* strain was created by PCR-amplifying 1kb homology arms up- and downstream of *bfmS* using NRA28 as template and cloning each in pUC18, creating pNRE31 and pNRE33, respectively. The two homology arms were subcloned by three-way ligation in pJB4648, creating the in-frame Δ*bfmS* allele (pNRE34). pNRE34 was recombined in NRA28 and allele exchange performed as above, resulting in strain NRA49.

To construct plasmids for *bfmR* overexpression, the *bfmR* gene of *A. baumannii* 17978 was amplified, cloned in pUC18 (creating pJE83), and subcloned in pEGE305 using EcoRI and PstI sites, creating pJE86. A mutant derivative harboring *bfmR*(D58A) was constructed by inverse PCR with bfmR-D58A-F, bfmR-D58x-R, and pJE83. The product was phosphorylated and circularized by ligation, creating pNRE137. The *bfmR*(D58A) allele was subcloned in EcoRI and PstI sites of pEGE305, creating pNRE138. To construct a *pilM*p-GFP reporter plasmid, the promoter region of *pilM* was PCR-amplified, cloned in pUC18, and subcloned in pEGE245 using SacI and KpnI. To construct plasmids for BfmR purification, *bfmR* or *bfmR*(D58A) was amplified, cloned in pUC18, and subcloned in pET28a via NdeI and BamHI sites to attach a cleavable N-terminal 6xHis tag, creating pNRE85 and pNRE127, respectively. *E. coli* BL21 (DE3) pLysS was used as host.

### Phos-tag and western blot analysis

Cells were grown to optical density (OD) ∼1, and 1 OD unit was harvested and lysed in 55ul BugBuster+0.1% Lysonase followed by addition of 18µl 4X SDS sample buffer (200mM Tris-HCl pH 6.8, 8% SDS, 0.4% Bromophenol blue, 40% Glycerol, 710mM BME). Samples were electrophoresed in 12% acrylamide gels containing Phos-Tag (50µM) and manganese (100mM). Gels were chelated in Transfer Buffer (50mM Tris, 380mM Glycine, 0.1% SDS, 20% Methanol) with 1mM EDTA, washed in Transfer Buffer without EDTA, and proteins transferred to low-fluorescence PVDF membranes. Total protein was detected with SYPRO Ruby blot stain (Invitrogen) following manufacturer’s protocol. Blots were blocked (5% milk in TBST) before incubation with rabbit anti-BfmR primary antiserum (Bai et al. 2023) (1:25,000 or 1:50,000), followed by goat anti-rabbit HRP conjugated secondary antibody (Invitrogen) (1:5,000). Blots were developed with Western Lightning Plus (Perkin Elmer) and imaged with a Chemi-Doc MP. Bands were quantified using ImageLab.

### Antibiotic resistance assay

Antibiotic resistance was assessed by measuring colony formation efficiency (6). Strains were grown to early post-exponential phase, resuspended in PBS, and serially diluted. Bacteria were spotted onto LB agar containing graded concentrations of antibiotic, followed by incubation at 37°C for 16 hours. Colony formation efficiency was calculated by dividing colony counts on antibiotic medium by colonies that formed on control medium lacking antibiotics. The minimum inhibitory concentration (MIC) is defined as the antibiotic concentration at which colony formation efficiency drops below 10^-3^.

### RT-qPCR

RNA was extracted with (RNeasy kit, Qiagen), DNase treated (Turbo DNase, Invitrogen), and reverse transcribed (SuperScript II, Invitrogen) using random primers. cDNA was amplified using gene-specific primers and PowerUp SYBR Green Master Mix (Invitrogen) on a QuantStudio3 (Applied Biosystems) in Fast Mode. Primers were validated to have an efficiency >90% by testing with serial dilutions of sheared gDNA template. Relative expression was quantified using 2^-ΔΔCt^ with *rpoC* as endogenous reference.

### Transcriptional reporter analysis

Strains harboring promoter fusions to GFP were cultured to early post-exponential phase, and 1 OD unit of cells was harvested and resuspended in 1 mL PBS. 200 µl of cells or PBS blank control was transferred into a black-walled, clear bottom 96 well plate (Corning). Fluorescence (ex480/em520nm) and OD (600nm) were read in a Synergy H1 microplate reader (Biotek). Fluorescence intensity was normalized by dividing by the *A*_600_ reading for each well.

### Animal experiments

Animal experiments were conducted in accordance with protocols approved by the Northeastern University Institutional Animal Care and Use Committee. Female C57BL/6 mice, 8-11 weeks of age, from Jackson Laboratories were used for all studies. JBA202 and NRA486 were cultured to early post-exponential phase, centrifuged, and resuspended in PBS. Mice were intraperitoneally injected with 100 µl of each strain suspension (∼5 x 10^6^ CFU). In survival studies, mice were monitored and those showing development of moribund state were euthanized. In experiments measuring bacterial burden, mice were euthanized 12 hours post-infection; blood was collected by cardiac puncture and transferred to K3-EDTA microtubes, and spleens were removed and homogenized in 1ml PBS. Samples were serially diluted and plated onto LB agar to enumerate CFUs.

### Purification of BfmR

NRE85 or NRE127 were cultured overnight in LB with Cm (25µg/mL) and Km (50µg/mL), diluted 1:100 into 1L of LB containing the same drug concentrations, and grown at 37°C until OD 0.6-0.8. 1mM IPTG was added and cultures were incubated for 4hrs at 30°C with shaking at 150rpm. Cells were harvested by centrifugation (4,000 x g for 10min) and stored at –80°C. Cell pellets were resuspended in cold Buffer A-5 (20mM Tris-HCl, pH 7.9, 500mM NaCl, 5mM Imidazole) with an EDTA-free protease inhibitor tablet (Roche) and kept on ice. Cells were lysed using a French press (Starstedt) or a Microfluidizer (Microfluidics Inc) and then centrifuged at 16,000 x g for 1hr at 4°C. Ni-NTA resin (Thermo Fisher) was added to a gravity flow column (Bio-Rad) in a cold room. The resin was equilibrated in 5CV cold Buffer A-5. The supernatant was then transferred to the column and allowed to flow through. The column was washed 3 times with 4CV of cold Buffer A-25 (20mM Tris-HCl, pH 7.9, 500mM NaCl, 25mM Imidazole). The protein was eluted from the column in 4CV of cold Buffer E (20mM Tris-HCl, pH 7.9, 500mM NaCl, 200mM Imidazole), transferred to 10kDa MWCO Float-a-Lyzers (Spectrum Labs), and dialyzed in Dialysis Buffer (20mM Tris-HCl, pH 7.5, 200mM NaCl). The protein was concentrated in 10kDa MWCO centrifugal concentrators (Millipore).

BfmR was further purified and analyzed by SEC using an Agilent 1260 HPLC system with ChemStation. BfmR was prepared in mobile phase (20mM Tris pH 7.5, 200mM NaCl). A 30 cm x 7.8 mmID TSKgel G2000SWXL column coupled with a guard column (4 cm x 6 mm) (TOSOH Bioscience) was equilibrated with 10CV of mobile phase at a flow rate of 0.5 mL/min under ambient temperature. 20 µL of sample was injected onto the column to purify, monitoring at single wavelength l = 280 nm.

### *In vitro* phosphorylation

Ammonium hydrogen phosphoramidate (PA) was synthesized according to published methods (24). Phosphorylation reactions were carried out in Buffer S (50mM Tris-HCl pH 7.4, 100mM NaCl, 10mM MgCl2, 50mM PA, 125µM BfmR) or Buffer D (20mM HEPES pH 6.5, 50mM NaCl, 5mM MgCl2, 0.5mM DTT, 50mM PA, 125µM BfmR) for 1hr at room temperature in an end-over-end rotator. Phosphorylation efficiency was analyzed on a 12% acrylamide Phos-Tag gel stained with SYPRO Orange (Invitrogen) following manufacturer’s instructions and visualized on a Chemi-Doc MP (BioRad).

### Chromatin immunoprecipitation (ChIP)

*A. baumannii* strains (NRA28, NRA29, and NRA49) were grown in biological triplicate to OD 0.5. Cultures were cross-linked by adding formaldehyde to 1% and rocking at room temperature for 30min. Reactions were quenched by adding Glycine to 250mM and rocking at room temperature for 15min. Cells were centrifuged at 5,000 x g for 1min, washed three times in 1mL PBS, and stored as pellets at −80°C. Cell pellets were resuspended in 500mL TES (10mM Tris-HCl, pH 7.5, 1mM EDTA, 100mM NaCl) and incubated with 35,000U of Ready-Lyse (Lucigen) for 10min at room temperature. 500mL of Buffer 1 (20mM KHEPES pH 7.9, 50mM KCl, 10% Glycerol) with a protease inhibitor tablet (Roche) was added and samples were subject to ultrasonication in a cup horn sonifier (Branson) with recirculating coolant for 7min (2min on, 2min rest, 50% output, 70% duty). Following sonication, samples were put on ice and then centrifuged at 13,000 x g for 20min at 4°C. Anti-FLAG magnetic beads (Sigma) were washed with 1mL IPP150 Buffer (10mM Tris-HCl pH 8, 150mM NaCl, 0.1% NP40) and resuspended in IPP150 Buffer. The buffer of the lysates was equilibrated to IPP150 Buffer by adding 5µL of 1M Tris-HCl pH 8, 20µL of 5M NaCl, and 10µL of 10% NP40, all per 1mL recovered supernatant. A 50µL aliquot of each lysate was taken for the “Input” control and frozen at −20°C. The supernatant was then added to 25µl of anti-FLAG beads and rotated end-over-end overnight at 4°C. The beads were washed by five rounds of magnetic separation, resuspension in 1mL IPP150 Buffer, and 2min of end-over-end rotation. The beads were then washed two times with 1mL TE (10mM Tris-HCl pH 7.5, 1mM EDTA) and 2min of end-over-end rotation. DNA-protein complexes were eluted from the beads by adding 150µL of Elution Buffer (50mM Tris-HCl pH 8, 10mM EDTA) and incubating at 65°C for 15min, inverting the tube every 5min. The supernatant was removed from the tube and stored. The beads were resuspended in 100µL of TE + 1% SDS and incubated at 65°C for an additional 5min. The supernatant was removed and pooled with the first elution. The crosslinking was then reversed for all eluted samples and the input samples by incubation at 65°C overnight. Samples were purified by using a PCR Purification Kit (Qiagen), eluting in 0.1X Elution Buffer. DNA was quantified by using the QuantiFluor dsDNA System (Promega). ChIP was confirmed by qPCR using primers specific for known BfmR-bound DNA compared to negative controls.

### ChIP-seq library prep, sequencing, and analysis

DNA sequencing libraries were prepared from ChIP-seq and Input DNA samples using the NEBNext Ultra II DNA Library Prep Kit for Illumina (New England Biolabs) following manufacturer’s protocol. After end prep and adapter ligation (with adaptors diluted 1:15), samples were amplified by 9 cycles of PCR without the preceding size selection step. Libraries were multiplexed by combining 25ng of each ChIP sample and 10ng of input control. Multiplexed libraries were size selected (Pippin HT, 175-600bp) and sequenced at the Tufts University Genomics Core Facility on an Illumina HiSeq 2500 (single-end 50bp reads), resulting in 7-27 million reads per library.

Fastx_clipper was used to trim adapter sequences and filter out low-quality reads from raw FASTQ reads (-v −l 20 -d 0 -Q 33 -a CTGTCTCTTATACACATCTCCGAGCCCACGAGAC). Filtered reads were aligned to the reference genome using bowtie (-q -m 1 -n 1 --best -y -S). Genome coverage was calculated using bamCoverage with CPM normalization. After quality assessment of mapped reads, analysis proceeded with all replicate samples (n=3) except for NRA28 (n=1) due to mapping anomalies indicating an incorrect source sample for 2 of its replicates. Prior to peak calling, each BAM file was downsampled to 500,000 reads using samtools to yield approximately 1x genome coverage. To identify genome regions enriched with aligned reads (peaks) after ChIP, peak calling was performed using MACS2 individually with each replicate ChIP sample, with NRA28 input sample as the control. Bedtools was used to identify common peak intervals present in at least two replicates. For these intervals, a custom script identified corresponding peaks in the replicates using the midpoint of the start and end coordinates. For each set of corresponding peaks, the script additionally calculated an average summit position (average of 0-based offsets from chromStart) and average fold enrichment (FE) value across replicates.

### ChIP-qPCR

Independent ChIP samples were isolated as above. Input samples were diluted to 0.04ng/µL and ChIP DNA was used at 1:10 dilution. Primers were designed to amplify within the ChIP-Seq peak intervals. Primers were validated to have efficiency > 90% by testing with a serial dilution series of sheared gDNA as template. Primers amplifying within *dnaA* (non-enriched in ChIP-seq data) was used as negative control. qPCR was carried out as described above for RT-qPCR. FE was calculated as 2^ΔΔCt^, where ΔΔCt = ΔCt(ChIP_control_-ChIP_target_)-ΔCt(Input_control_-Input_target_).

### Microscale thermophoresis (MST)

Purified BfmR (without or with phosphorylation as above) was serially 2-fold diluted and combined with duplex DNA (5’ 6-FAM-labeled on one strand) at 20nM in Buffer D. After incubation for 10 minutes at room temperature, mixtures were loaded in technical duplicate into Premium Capillary Chips (Nanotemper). Interactions were measured on a Nanotemper Monolith NT Automatic instrument via Mo.Control software in Expert Mode (Nano-BLUE excitation mode, 85% excitation power, Medium MST power). An average normalized fluorescence (F_norm_) value was calculated from technical duplicates. Replicate capillaries with F_norm_ outliers or MST trace abnormalities indicative of aggregates were excluded from analysis. ΔF_norm_ was calculated by subtracting the average F_norm_ value from the Mo.Affinity-generated “Unbound” value. ΔF_norm_ was plotted vs BfmR concentration and Kd calculated from non-linear regression analysis (Specific binding with Hill slope) using Graphpad Prism 10.

### Identification of direct target ORFs and pathway enrichment

Direct BfmR target genes were compiled using the criteria detailed in the main text. To identify ORFs with significant differential gene expression in *bfmRS* mutants in published studies, RNA-seq data were compiled from the supplemental datasets in (6) (17978) and (17) (17961). Corresponding loci between the strains were determined by identifying positional orthologs with progressiveMauve (25). Genes forming transcriptional units were determined from the BioCyc *A. baumannii* ATCC 17978 database (26,27). Overrepresented Gene Ontology (GO) terms among sets of directly activated or repressed genes were identified using the FUNAGE-Pro web server using “Auto detect threshold” (28).

### sRNA analysis

The sRNA loci previously identified in ATCC 17978 by Kroger et al (29) were compiled in a .gtf file via a custom Python script. Sequences were extracted and BLAST+ (30) was used to identify corresponding regions in ATCC 17961 (GenBank accession CP065432.1). BLAST+ hits were filtered for at least 80% sequence identity and at least 99% of the sRNA’s length in 17978, and used to generate a .gtf file for ATCC 17961 via Python script. To re-analyze published RNA-seq data for differential expression of sRNA genes, RNA-seq reads (Single-end 50) for *A. baumannii* 17978 WT, Δ*bfmR*, Δ*bfmS*, and Δ*bfmRS* in exponential phase (6) were quality-checked (FastQC) and aligned to NZ_CP012004.1 (Bowtie 2.5.2 with -- local --very-sensitive-local settings (29,31). Alignments were converted to .bam format and sorted using samtools 1.19.2 (32). Aligned reads were assigned to sRNA features using the featureCounts program in the Subread 2.0.6 package (33) with appropriate settings for single-end reads. Differential expression analysis was performed using DESeq2 1.42.0 (34) with LFC shrinkage performed via the apeglm package (35). sRNA features were filtered to include only features with raw read counts ≥10 in at least 3 samples during DESeq2 analysis. Alignment statistics were inspected using MultiQC (36). Paired-end Illumina sequencing reads generated by Palethorpe et al using ATCC 17961 (17) were aligned to CP065432.1 using --local --very-sensitive-local settings. The ATCC 17961 .gtf file described above was used to specify features for featureCounts, and appropriate settings were used for paired-end reads. Read quality inspection, alignment, .bam file processing, aggregation of reads to genomic features, and differential expression analysis were performed as above.

## RESULTS

### Phosphorylation of BfmR depends on Asp58 and is modulated by BfmS *in vivo*

To examine the importance of BfmR phosphorylation *in vivo*, we introduced a chromosomal *bfmR*(D58A) mutation into *A. baumannii* 17978 by allelic exchange and used phosphate-affinity gel electrophoresis to measure levels of phospho-BfmR (BfmR∼P; Materials and methods) (9). Consistent with previous work, WT control bacteria have two forms of BfmR, with a fraction of about 20% showing a migration shift indicating BfmR∼P (Fig. 1A). Bacteria lacking BfmS show an increased proportion of BfmR∼P (Fig. 1A), as observed previously (9,17). Replacing BfmR with the D58A variant completely abolished the BfmR∼P band in both BfmS^+^ and Δ*bfmS* backgrounds (Fig 1A). These results agree with the prediction that Asp58 is the key, single site of phosphorylation in BfmR.

**Fig. 1.**
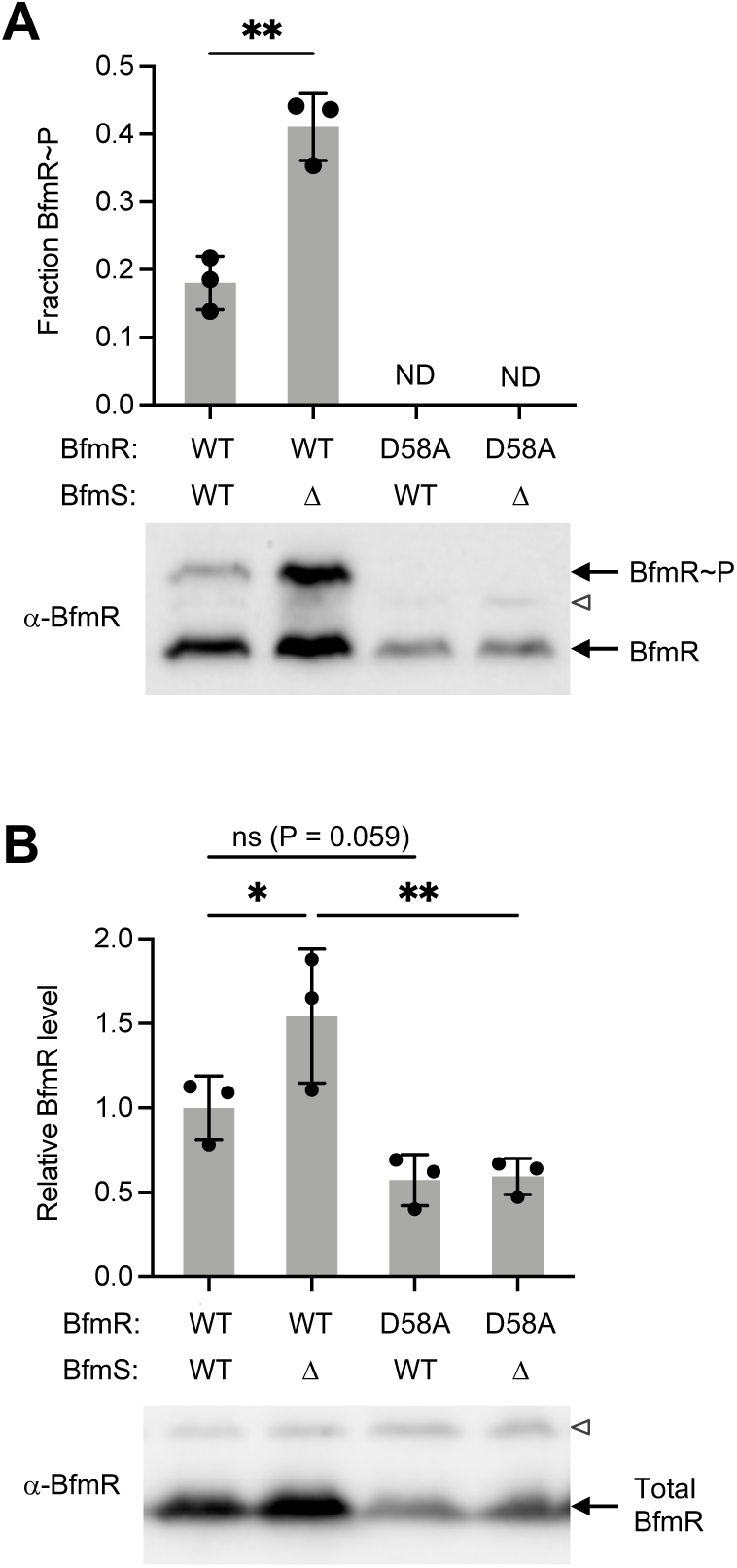
D58A mutation blocks BfmR phosphorylation. (A) Phos-tag immunoblot analysis of BfmR phosphorylation using BfmR antiserum. Graph shows fraction BfmR∼P/total BfmR (mean ± s.d., n=3). Results were analyzed by unpaired t-test. ND, no visible BfmR∼P band detected. (B) Immunoblot analysis of total BfmR levels using separate gels without phos-tag reagent. Data show mean ± s.d., n=3. P values shown from the indicated comparisons after one-way ANOVA with uncorrected Fisher’s LSD test. Representative blots are shown below each graph; open arrowhead denotes non-specific protein band reacting weakly with BfmR antiserum. ns, P>0.05; *, P≤0.05; **, P≤0.01.

Compared to the WT strain, overall BfmR protein levels appeared to increase in Δ*bfmS* (as noted previously (9)) and decrease in both *bfmR*(D58A) mutants. To measure changes in BfmR further, we analyzed the same lysates by SDS-PAGE without the phosphate affinity reagent. The resulting Western blots revealed that the Δ*bfmS* strain had an increase in the total amount of BfmR protein by approximately 1.5-fold, and the D58A strains had a decrease by a similar magnitude (Fig. 1B, S1A). These results are consistent with positive autoregulation at the *bfmRS* locus.

### BfmR requires phosphorylation to regulate gene expression and promote resistance and virulence

To test the prediction that phosphorylation is essential to BfmR activity, we measured the effects of *bfmR*(D58A) on key BfmR-dependent phenotypes, comparing the mutant to both isogenic WT and a complete *bfmRS* knockout (Δ*bfmRS*). First, we examined gene expression at two loci, *slt* and *pilM*, identified previously by RNA-seq to be under strong positive and negative control by BfmRS, respectively (6,17). RT-qPCR showed that *slt* expression was decreased and *pilM* expression was increased in the Δ*bfmRS* mutant compared to the WT, as predicted by RNA-seq, and the *bfmR*(D58A) mutant showed almost exactly the same effects on expression for both genes (Fig. 2A). A similar result was seen when testing transcriptional GFP reporter fusions to the *slt* and *pilM* promoters. Compared to WT, the Δ*bfmRS* and *bfmR*(D58A) mutations both caused promoter activity of *slt* to decrease and *pilM* to increase, by magnitudes indistinguishable between the two mutants (Fig. 2B). Second, we tested the effects of *bfmR*(D58A) on resistance to two antibiotics, mecillinam and rifampicin. As shown previously, the Δ*bfmRS* mutation causes major defects in the ability of bacteria to grow on medium containing these antibiotics, lowering the mecillinam MIC from 64 µg/ml (with WT) to 4 µg/ml, and the rifampicin MIC from 2 µg/ml (with WT) to 0.5 µg/ml (Fig. 2C). The *bfmR*(D58A) mutant behaved identically to the Δ*bfmRS* knockout in these resistance assays (Fig. 2C). Together, these results argue that BfmR requires phosphorylation for its ability to modulate gene expression and confer intrinsic antibiotic resistance.

**Fig. 2.**
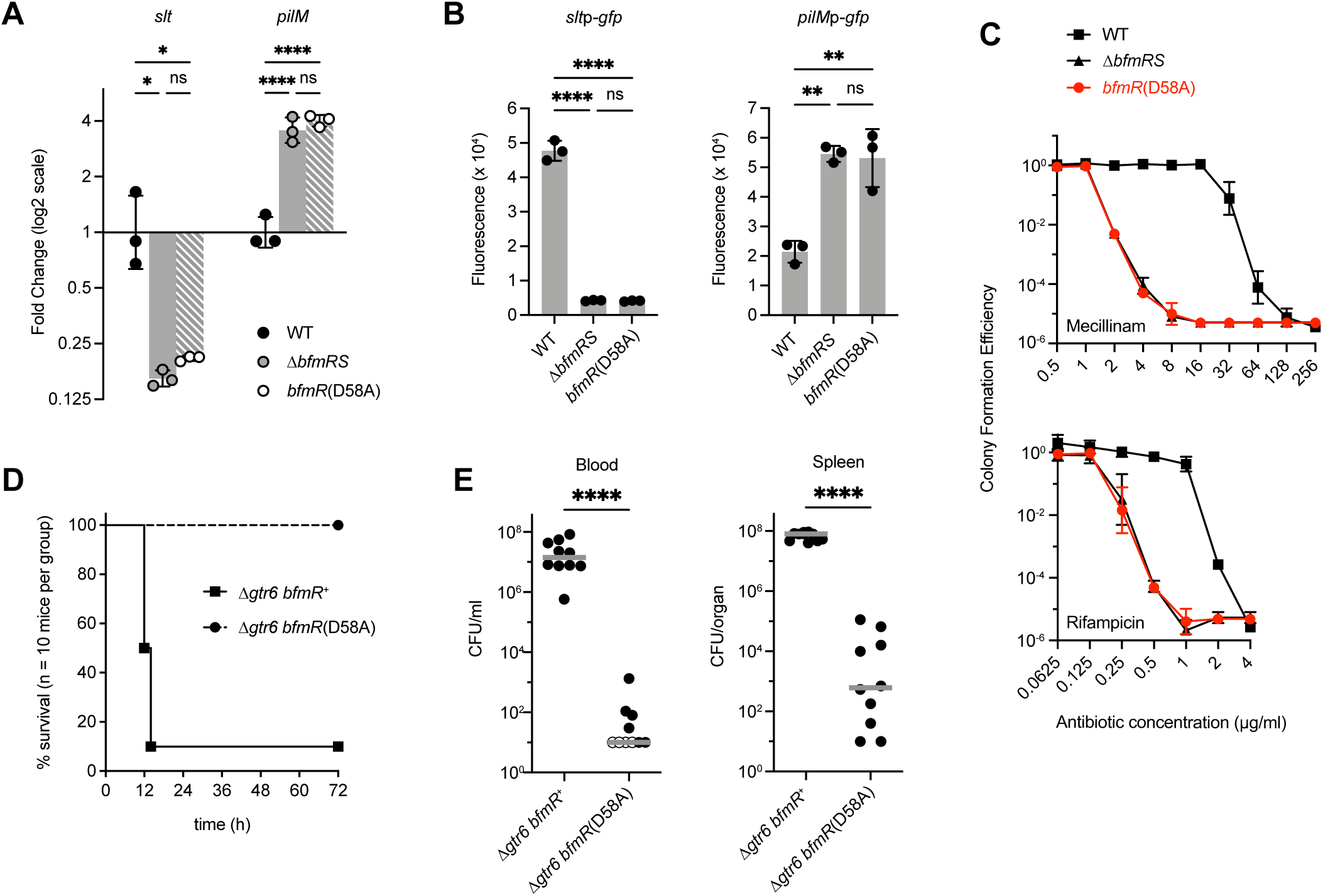
*bfmR*(D58A) mutant shows BfmRS-null phenotypes. (A) Analysis of *slt* and *pilM* transcription in the indicated *A. baumannii* strain by RT-qPCR. Data show geometric mean ± s.d. (n=3). (B) Analysis of transcriptional reporter fusions to *slt* and *pilM* promoters. Bars show mean normalized fluorescence (fluorescence units/A_600_) ± s.d. (n=3). (C) Antibiotic resistance assays. Colony forming efficiency was measured on solid medium with graded doses of mecillinam (top) or rifampicin (bottom) compared to control medium lacking drug. Data points show geometric mean ± s.d. (n=3). (D-E) Results from the mouse sepsis model. D shows mouse survival after intraperitoneal injection of bacteria (mean dose 7.3×10^6^ CFU with Δ*gtr6 bfmR*^+^, 5.4×10^6^ CFU with Δ*gtr6 bfmR*(D58A). Results were pooled from two independent experiments with 5 mice per group in each experiment. Survival curves are significantly different (Gehan-Breslow-Wilcoxon Test, p<0.0001). E shows bacterial burdens in the blood and spleen 12 hours after infection. Mean CFU in the inocula were 3.9×10^6^ [Δ*gtr6 bfmR*^+^] and 4.0×10^6^ [Δ*gtr6 bfmR*(D58A)]. Results were pooled from 2 independent experiments with 5 mice per group in each experiment. Each symbol represents one animal. Open symbols denote value below the limit of detection. Gray lines are medians. Differences between mutant and control were evaluated by Mann Whitney test. ns, P>0.05; *,P≤0.05; **, P≤0.01; ****, P≤0.0001.

Finally, we tested the effect of *bfmR*(D58A) mutation on virulence in a mouse septicemia model. Because strain 17978 shows low virulence, to establish a bloodstream infection we took advantage of the Δ*gtr6* derivative of 17978, in which the capsule is modified to a form found naturally to confer strains with high virulence (37). A Δ*gtr6 bfmR*(D58A) double mutant was constructed and compared with its *bfmR*^+^ parent in the mouse model by injection of ∼5 x 10^6^ bacteria intraperitoneally. Infection with the Δ*gtr6 bfmR*^+^ control bacteria led to lethal disease in 90% of mice (Fig. 2D) and high bacterial loads in the bloodstream and spleen (Fig. 2E). By contrast, the isogenic Δ*gtr6 bfmR*(D58A) mutant was unable to cause lethality and showed dramatic 6 and 5-log decreases in bacterial burdens in the bloodstream and spleen, respectively, compared to the control (Fig. 2D,E). These data demonstrate the essential role of BfmR Asp58 in the virulence of *A. baumannii*.

Based on analogy with other OmpR/PhoB systems (38), loss of phosphorylation is the likely defect responsible for the above null phenotypes of *bfmR*(D58A); nevertheless, we considered the alternative possibilities that low BfmR protein levels (Fig. 1B) or largescale structural changes were contributing factors. To rule out the first possibility, we assessed the degree to which raising BfmR^D58A^ protein production to WT levels could restore activity of the regulator. To this end, IPTG-dependent *bfmR*(WT) or *bfmR*(D58A) alleles were introduced into a Δ*bfmR* strain via plasmid and induced with an IPTG concentration (25 µM) yielding BfmR levels just above that seen in a WT strain with native *bfmR* (Fig. 3A, S1B), thus correcting for the low protein levels associated with autoregulation. Despite the higher protein level, the D58A variant, which remained completely phosphorylation deficient (Fig. S1C), was unable to confer any increase in mecillinam or rifampicin resistance when compared to the Δ*bfmR* parent strain harboring the vector alone (Fig. 3B). By contrast, at the same level of induction WT BfmR was able to increase resistance to the same antibiotics dramatically, reaching a magnitude equivalent to or greater than that seen with the control WT strain (Fig. 3B). The increased mecillinam resistance conferred by overexpressed *bfmR*(WT) likely reflects the higher amount of BfmR∼P in this strain compared to the WT control (Fig. S1C), an effect also seen with the Δ*bfmS* mutant (see (6) and Fig. S3B). To address the second possibility, we examined whether the D58A mutation could exert dominant negative effects in a BfmR^+^ background. The same IPTG-dependent *bfmR* alleles were introduced into WT *A. baumannii* and induced with an IPTG concentration (75 µM) yielding 8-9-fold higher expression over native levels (Fig. 3C, S1D). Growth was assessed on solid medium containing antibiotics at sub-MIC. Unlike the WT allele which increased growth on mecillinam medium, overexpression of *bfmR*(D58A) caused a significant decrease in growth compared to the vector-only control (Fig. 3D, left). Overexpressing *bfmR*(D58A) caused a similar decrease in growth on medium containing rifampicin, whereas similarly expressed *bfmR*(WT) had no effect (Fig. 3D, right). The dominant-negative ability of BfmR^D58A^ argues against wholesale structural deformities in the protein, as it must be in a form competent for interactions, likely weak, that interfere with BfmR function. Together, these results support the idea that lack of phosphorylation, rather than lowered protein levels or structural instability, are the root of the dramatic antibiotic resistance and virulence defects seen with the BfmR^D58A^ variant.

**Fig. 3.**
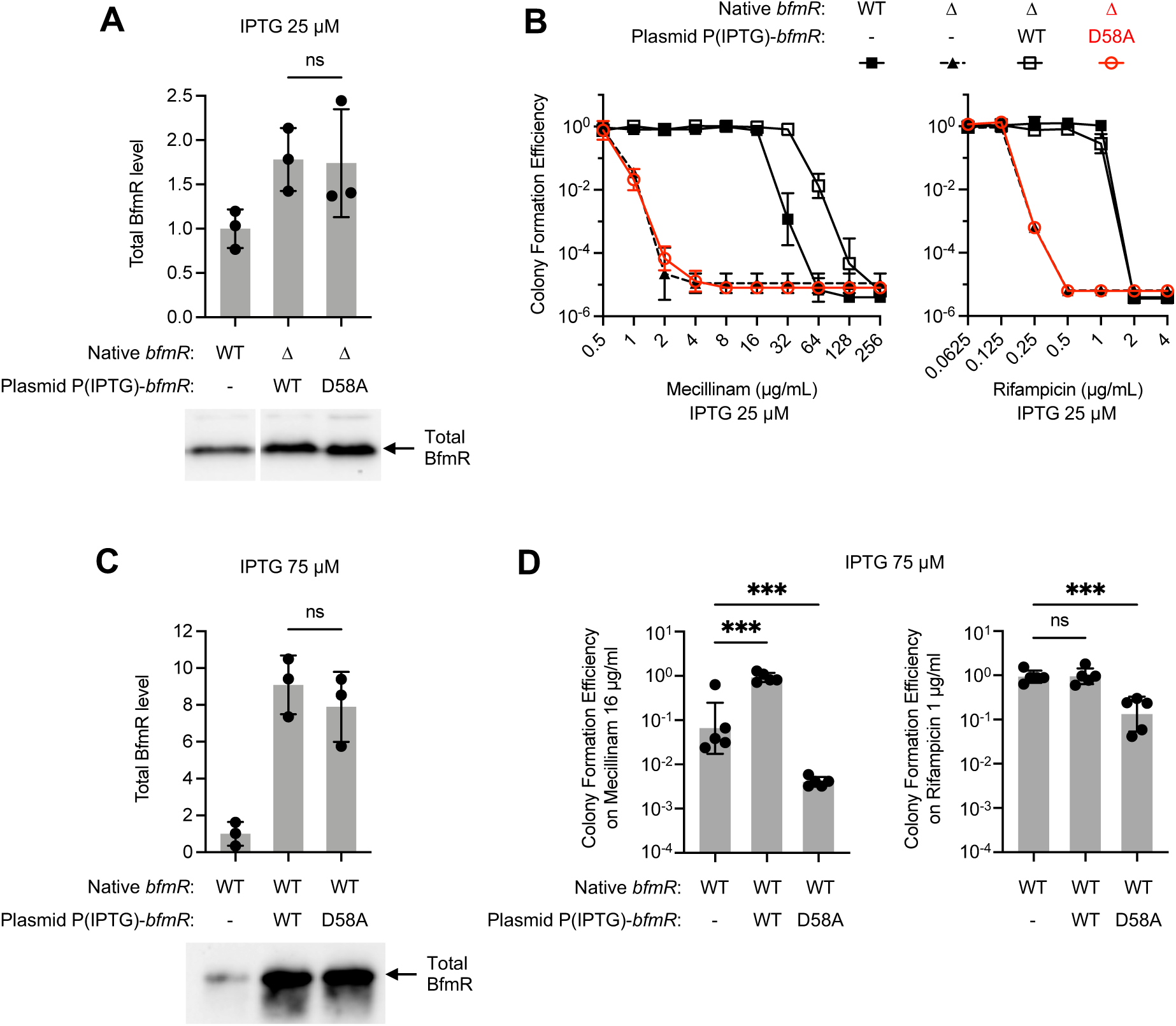
When overexpressed, the phosphorylation-deficient *bfmR*(D58A) allele lacks complementing ability but has dominant negative effects. Strains harbored the indicated *bfmR* allele in its native chromosomal locus or via plasmid. Expression from plasmid-borne *bfmR* was driven by 25 (A, B) or 75 (C, D) µM IPTG. A, C. Immunoblot analysis of total BfmR levels. Data show mean ± s.d. (n=3). A representative blot is shown below each graph. B, D. Antibiotic resistance assays. Data show geometric mean ± s.d., n=3 (B), n=5 (D). In D, one-way ANOVA with Dunnett’s test was used to compare *bfmR*-overexpressed strains to the control (WT with empty vector). ns, P>0.05; ***, P≤0.001.

### Phosphorylation induces dimerization of BfmR

Typically, phosphorylation enhances dimerization of OmpR family response regulators (39), and we examined whether BfmR follows this paradigm. BfmR was shown previously to form dimers upon treatment with BeF_3_^-^(23). To analyze phosphorylation-induced dimerization, we purified BfmR and used the small phosphodonor PA to phosphorylate the protein *in vitro* (Materials and Methods). Phosphorylation by PA was highly efficient, causing almost all of the BfmR protein to exhibit a phosphate affinity gel shift, indicating the presence of BfmR∼P (Fig. 4A). Heating at 95°C abolished the shift, consistent with a heat labile phospho-aspartate bond in the PA-treated protein. By contrast with WT BfmR, PA treatment did not cause an analogous shift for purified BfmR^D58A^ (Fig. 4A), further evidence of the variant’s defect in phosphorylation. We next analyzed oligomerization by using SEC. We used a set of protein standards to determine approximate molecular weights of the states of BfmR, with the caveat that protein shape influences the accuracy of the estimated size values. Untreated WT BfmR (theoretical molecular weight of 27.4 kDa) was eluted in a uniform peak, corresponding to an apparent size of 40.7 kDa (Fig 4B, Fig. S2A). PA treatment caused almost all of the protein to elute earlier, at a higher apparent molecular weight (55.5 kDa) very close to the size expected for a homodimer (theoretical molecular weight of 55 kDa) (Fig. 4B, Fig. S2A). These results suggest that the unphosphorylated protein is a monomer, with an asymmetric, flexible shape (23) likely responsible for a large apparent Stokes radius during gel filtration, as observed previously with other response regulators (40,41). By contrast, phosphorylation appears to induce a dimeric form that behaves more like a globular particle under the same gel-filtration conditions. Notably, in contrast to the result with the WT protein, BfmR^D58A^ eluted at the monomer size but did not show any change in elution pattern with PA treatment, indicating that there were no dimers formed (Fig. 4B). These results lend further support for the model that BfmR is phosphorylated at Asp58 and that phosphorylation induces dimerization.

**Fig. 4.**
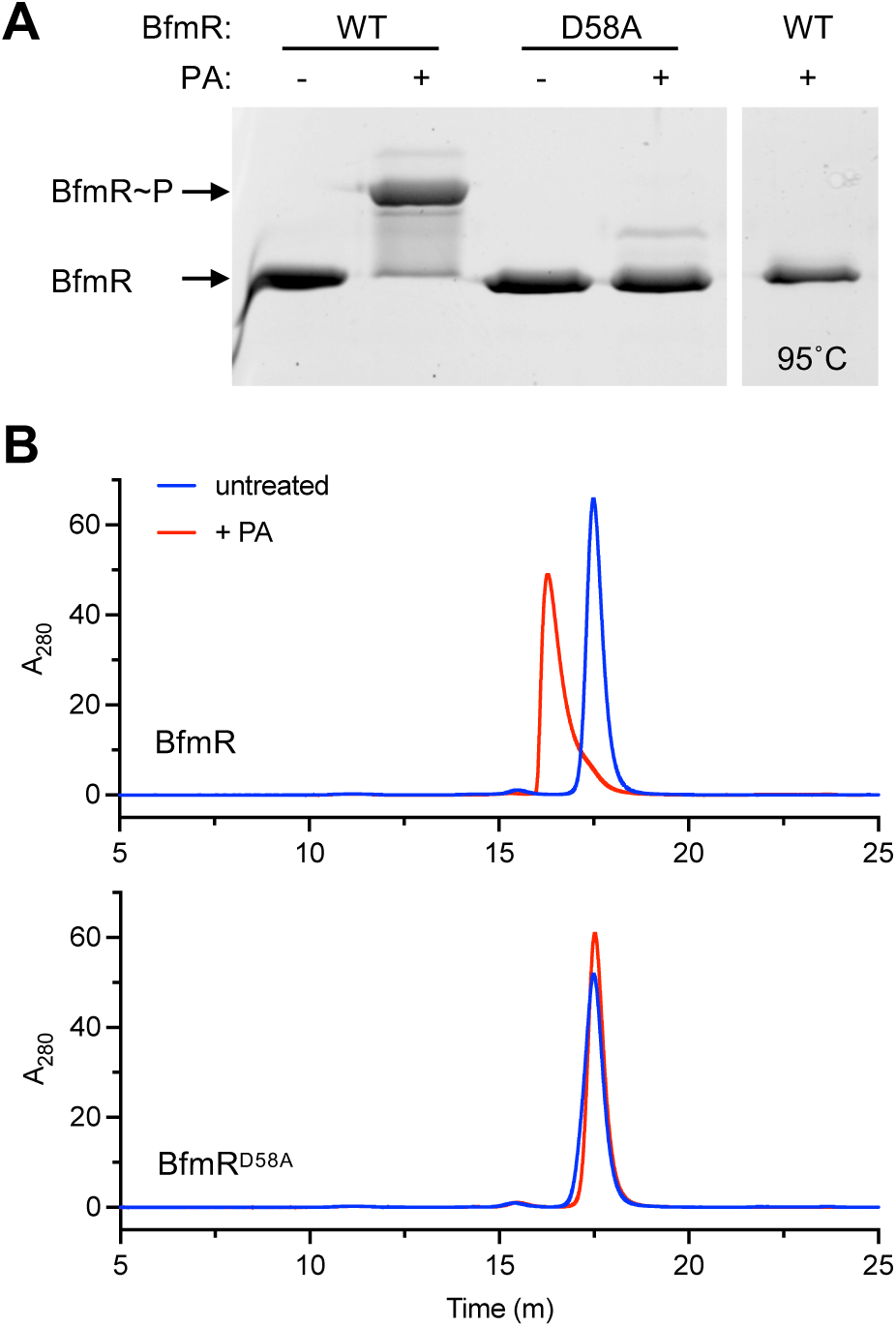
Phosphorylation of BfmR *in vitro* induces dimerization. (A) Phos-tag gel analysis of *in vitro* BfmR autophosphorylation. Purified BfmR protein was incubated with PA for one hour in Buffer S and combined with SDS loading buffer. In the last lane, the sample was heat inactivated at 95°C for 10min before addition of SDS buffer. Gel was stained with SYPRO orange. (B) Analysis of BfmR dimerization via analytical SEC. Purified BfmR or BfmR^D58A^ was incubated with PA at 50µM (red) or no PA (blue) in Buffer S. Representative HPLC chromatograms are shown (representative of n=3).

### BfmR binds to sites throughout the genome in phosphorylation-dependent fashion

BfmR has a large regulon but the extent of the DNA targets that the regulator directly binds is not known (6,17). To identify genome-wide sites of BfmR binding and the role of phosphorylation in these interactions, we performed ChIP analysis, taking advantage of three strain backgrounds with varying amounts of BfmR∼P: Δ*bfmS* (BfmR∼P high), WT (BfmR∼P low), and *bfmR*(D58A) (no BfmR∼P). To facilitate BfmR immunoprecipitation, we constructed versions of the above strains in which the native *bfmR* is fused to an epitope sequence (C-terminal 3XFLAG tag). Phos-tag gel analysis and antibiotic resistance assays revealed the expected pattern of BfmR phosphorylation and mecillinam resistance in the resulting strains (Fig. S3), confirming that the fusion tag preserves BfmR functions. BfmR-bound DNA from the above strains was subsequently fragmented, immunoprecipitated and detected by Illumina sequencing (Materials and Methods).

Alignment of the resulting sequence reads to the *A. baumannii* genome revealed numerous ChIP-seq signals distributed throughout the chromosome and plasmids in the strains producing BfmR∼P (Δ*bfmS* and WT), and a comparative dearth in *bfmR*(D58A) (Fig 5A). BfmR ChIP-seq peaks showing significant enrichment over the input control were determined, using a q-value cutoff of 0.05 (Materials and Methods). This approach resulted in identification of 248 distinct BfmR binding sites in the Δ*bfmS* strain, 105 in the WT, and only four in the *bfmR*(D58A) mutant (Dataset S1 and Fig. 5C). 10 and 8 of these peaks in Δ*bfmS* and WT, respectively, were present in plasmids (Dataset S1 and Fig. 5B). The vast majority of peaks detected in the WT (81/105) and all 4 peaks identified in *bfmR*(D58A) were at sites identical to those found in the Δ*bfmS* strain. This suggests that increasing the level of BfmR∼P has the general effect of increasing the range of loci bound by BfmR. Furthermore, most of the shared sites (66/81) showed higher FE in Δ*bfmS* compared to the WT or *bfmR*(D58A) (Fig. 5D, blue lines), although 15 sites had the highest FE in the WT strain (Fig. 5D, red lines). The Δ*bfmS* and WT strains each showed a large dynamic range in ChIP-seq FE values (1.5-11.8, and 1.7-6.6, respectively), likely reflecting heterogeneity among genome-wide BfmR-target DNA interactions. We identified the summits of the peaks and determined their position relative to the closest ORFs based on genome annotations (Dataset S1). Most of the BfmR binding sites were intergenic (54% and 69% of Δ*bfmS* and WT peaks, respectively) and were preferentially situated between 25 and 125bp upstream of the nearest ORF start (Fig. 5E). This biased distribution provides evidence that BfmR is interacting with promoter regions. Peaks within coding regions also tended to be close to an ORF start site, with a preference for occurring within 0-75bp from the corresponding start codons (Fig. 5E).

**Fig. 5.**
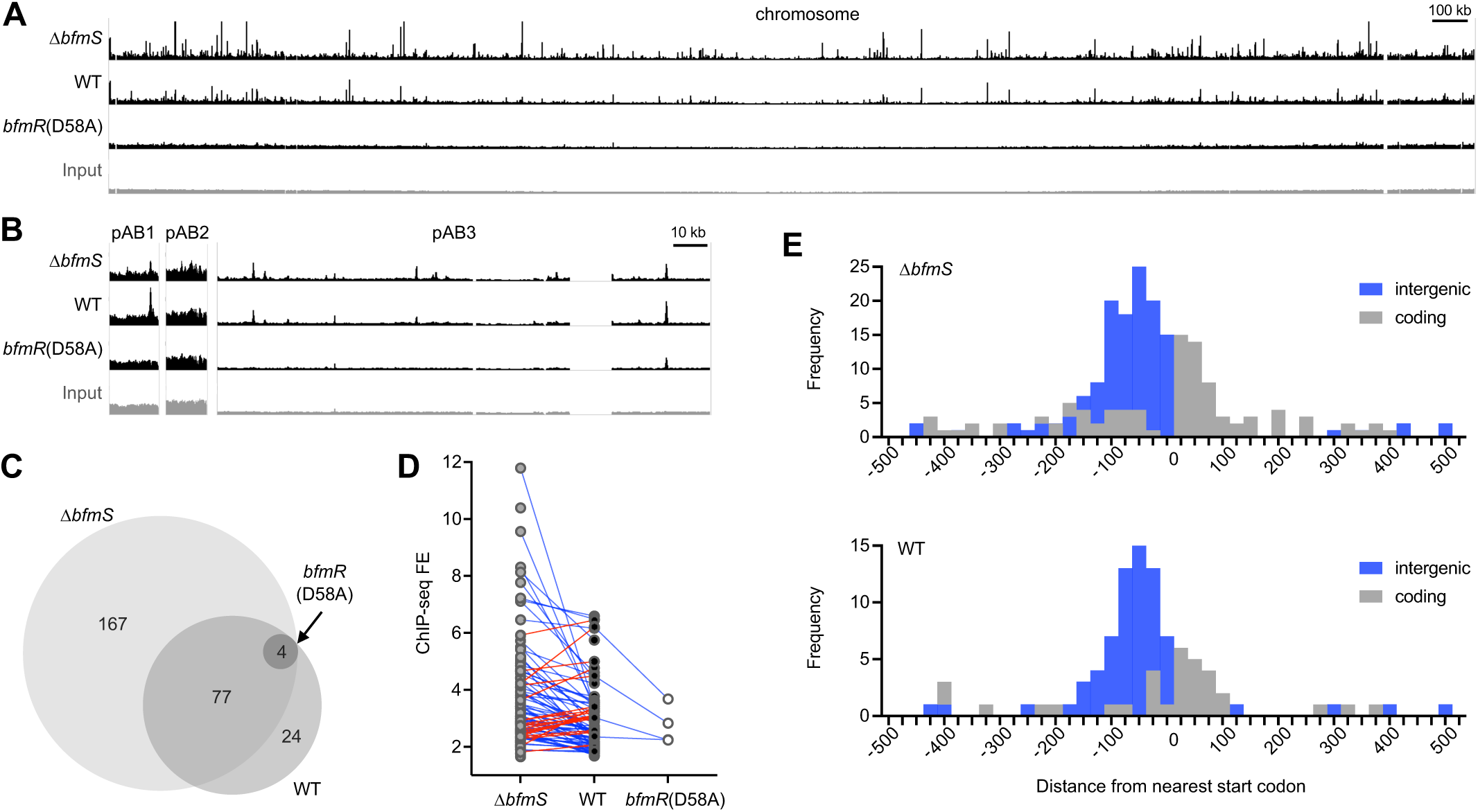
BfmR binds to sites throughout the *A. baumannii* genome in a manner dependent on phosphorylation. (A-B) Genome-wide BfmR binding sites revealed by ChIP-seq. Tracks show reads after ChIP-seq with the indicated strain, or input reads before ChIP from WT strain (input), mapped to the 17978 chromosome (A) or plasmids (B). Track height is 300 reads. Gap regions in chromosome and pAB3 read coverage correspond to unmapped endogenous transposon-related sequences. Δ*bfmS* refers to NRA49, WT refers to NRA28, and *bfmR*(D58A) refers to NRA29. (C) Venn diagram illustrating the number and relationships of BfmR binding sites detected in each strain. (D) Analysis of the relationship between FE for a binding site and strain background. Blue lines indicate sites in which FE was higher in Δ*bfmS* than WT. Red lines indicate sites in which FE was higher in WT vs Δ*bfmS*. (E) Histograms show the distance of the peak summit from the nearest ORF start codon. Binding sites in intergenic regions are shaded blue, and those within coding regions are shaded grey.

To validate the ChIP-seq findings, we performed independent ChIP-qPCR experiments using a separate set of bacterial cultures (Materials and Methods). qPCR probes were designed using the ChIP-seq peak information for 16 binding sites in predicted promoter regions, and BfmR binding was evaluated by examining ChIP-qPCR FE over input control. BfmR binding was again detected at all sites in strains producing BfmR∼P, while almost no sites showed binding in the *bfmR*(D58A) background (Fig. 6A). The FE values were higher in the Δ*bfmS* than the WT strain at all but one of these sites, consistent with the findings from ChIP-seq. With the *gidA* promoter, which was one of the few sites having a ChIP-seq peak in the *bfmR*(D58A) strain (Dataset S1), a binding signal was again detected in *bfmR*(D58A) by ChIP-qPCR (FE of 4.2), albeit at levels lower than that seen in strains producing BfmR∼P. Overall, the ChIP-seq and ChIP-qPCR FE values were positively correlated in both Δ*bfmS* and WT samples (Fig. 6B). Together, these results reveal the genome-wide binding sites of BfmR and support the notion that interaction with targets is increased with BfmR phosphorylation.

**Fig. 6.**
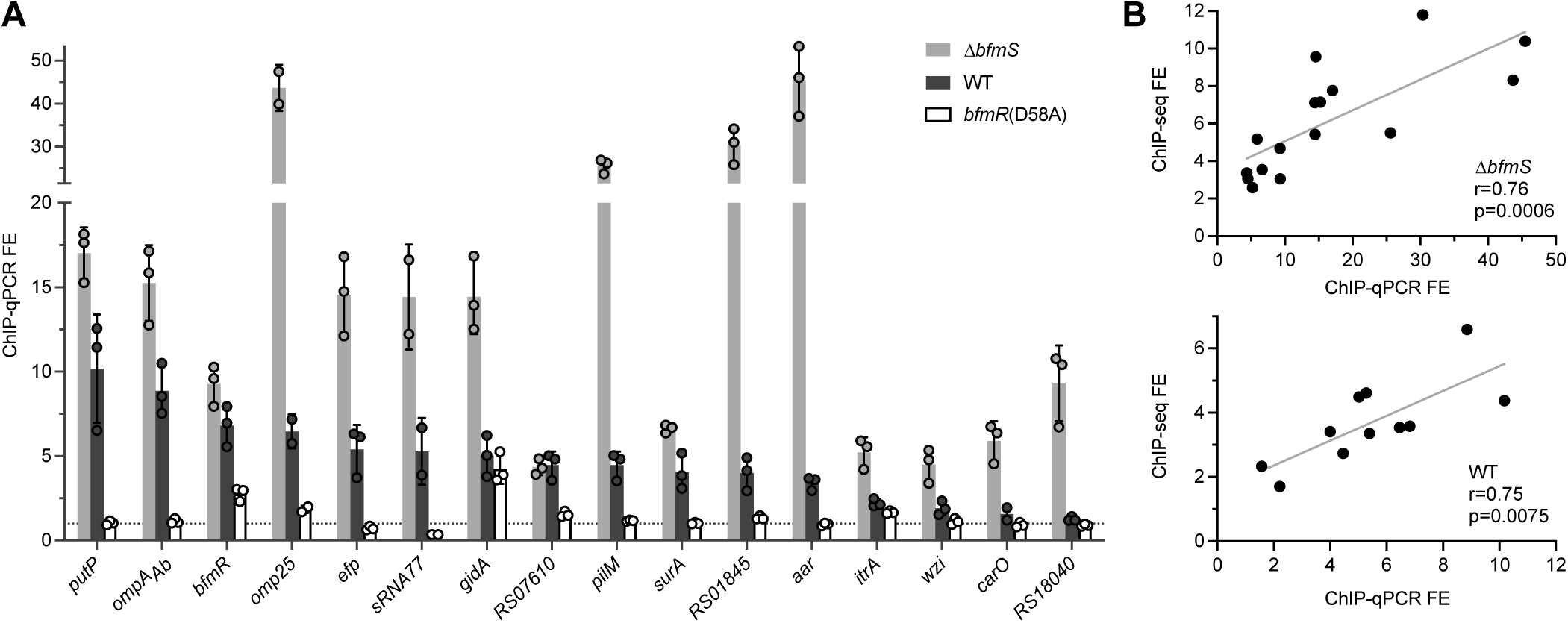
Independent ChIP-qPCR experiments validate phosphorylation-dependent BfmR binding at select target sites. (A) Independent ChIP was performed with NRA49 (Δ*bfmS*), NRA28 (WT), and NRA29 [*bfmR*(D58A)]. qPCR was used to calculate fold enrichments at binding sites predicted by ChIP-seq. Binding sites are labelled by the closest gene. Dotted line denotes FE value of 1. Data points show mean ± s.d. (n=3; except *omp25* and sRNA77, n=2). (B) Correlation of the FE values from ChIP-Seq and independent ChIP-qPCR for NRA49 (top) and NRA28 (bottom). Correlations were evaluated by Pearson’s r. Gray line shows simple linear regression.

### Identification of a BfmR binding motif

To determine a common DNA sequence motif recognized by BfmR∼P, we used the MEME-ChIP software suite (42–44). We focused on the subset of 87 BfmR ChIP-seq peaks in the Δ*bfmS* strain showing FE of 3 or greater, and searched sequences ±250bp from the summit of these peaks (Materials and Methods). The search identified two motifs: a 15bp sequence containing two direct repeats (of the consensus sequence “TTACA”) (E-value 1.3×10^-13^; Fig. 7A) and an 8bp sequence (E-value 3.9×10^-3^; Fig. 7B) that contains one of the repeats. Consistent with direct binding of BfmR to the DNA enriched in ChIP-seq, the motifs were preferentially located near the ChIP-seq peak summits (Fig. 7C). Interestingly, while the 8bp motif showed clear central enrichment, many of the 15bp motifs tended to be slightly offset from the peak center by about 30bp (Fig. 7C). We found that the vast majority of BfmR binding sites contained the motifs. For instance, a scan of the ChIP-seq sites in Δ*bfmS* for motifs using FIMO with p-value cutoff of 1 × 10^−3^ revealed that 88% and 81% of BfmR binding sites had at least one occurrence of the 15bp and 8bp motif, respectively (Dataset S2).

**Fig. 7.**
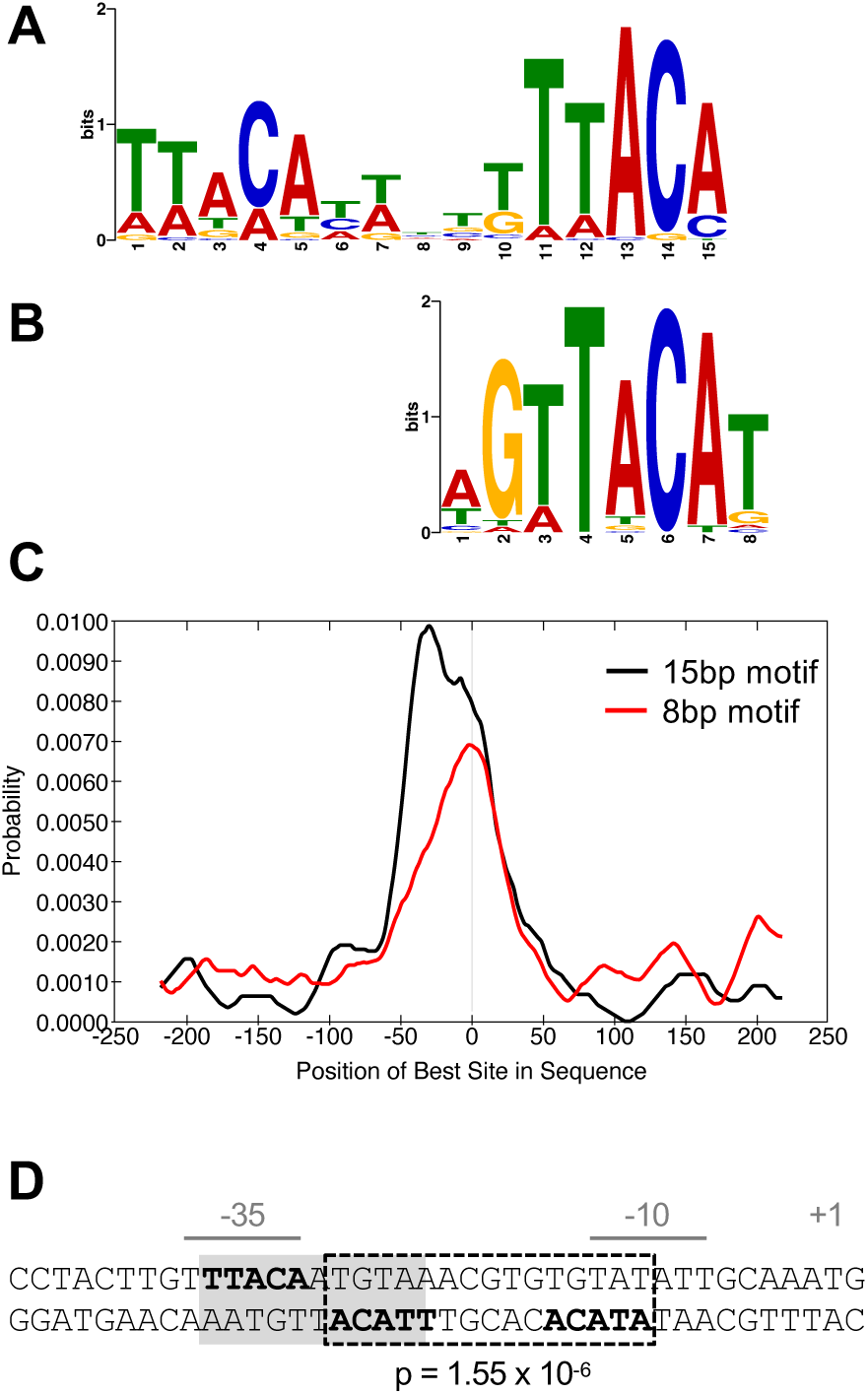
BfmR-targeted DNA motifs. (A-C) Identification and distribution of BfmR-binding motifs using a set of 87 input sequences associated with a ChIP-seq peak FE >3 in NRA49 (Δ*bfmS*). (A) 15bp BfmR-binding motif identified from MEME motif discovery program (E-value 1.3×10^-13^). (B) 8bp binding motif identified from STREME motif discovery program (E-value 3.9×10^-3^). (C) Distribution of the best matches to the motif in the binding site sequences as analyzed by using CentriMo. The vertical line in the center of the graph corresponds to the center of the sequences. (D) Location of a BfmR binding motif in the BfmR promoter. The promoter identified in (8) is shown. The repeated half sites are in bold. Dotted box denotes a 15bp direct-repeat BfmR binding motif determined by FIMO (p-value shown). Grey shading denotes the previously identified inverted repeat (23).

The TTACA consensus sequence for BfmR binding resembles parts of the recognition motifs identified for OmpR and RstA in *E. coli* (45–47). In addition, the TTACA sequence had been identified previously within the *bfmRS* promoter in the context of two inverted repeats separated by 1bp (Fig 7D, grey shading) (23). Of note, half of the 11bp inverted site overlaps with a highly significant direct repeat BfmR motif (p-value=1.55×10^-6^) that is located between the predicted −35 and −10 elements (Fig 7D, dotted box). The 15bp direct repeat sequence within the *bfmRS* promoter was part of a cluster of three 15bp motifs and five 8bp motifs in this region (Fig. 8B). The 11bp inverted repeat arrangement was not detected elsewhere in the *bfmR* promoter region and was in general less frequent across genome-wide BfmR binding sites than the direct repeat, occurring in only 35% using the same FIMO search criteria as above.

**Fig. 8.**
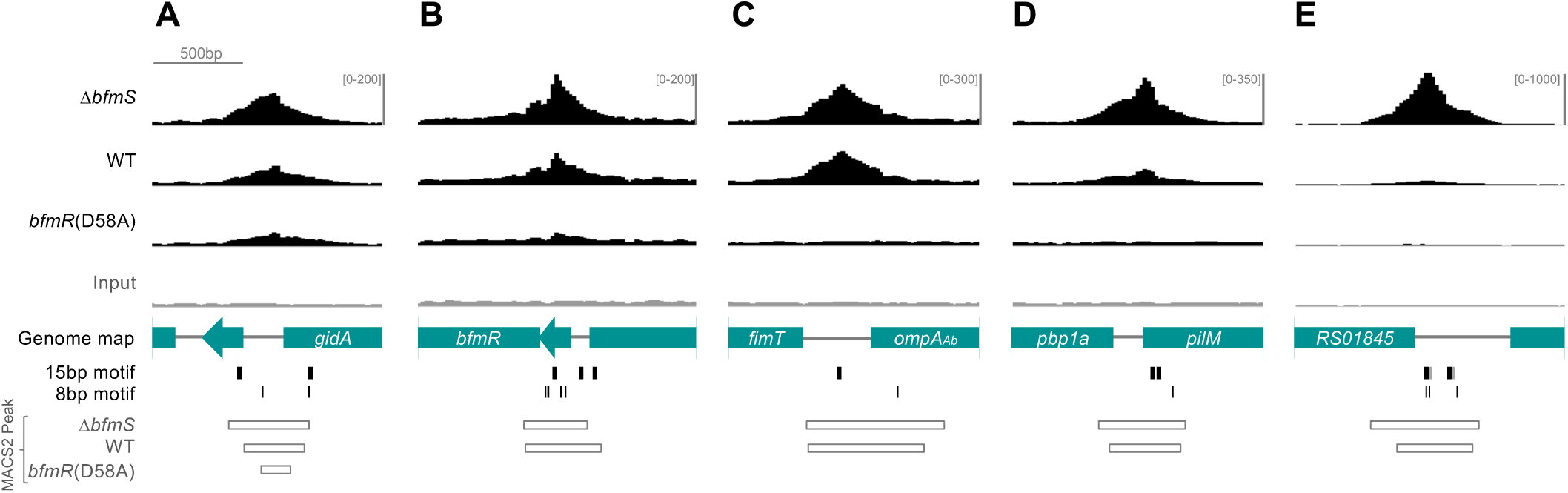
BfmR ChIP-seq peaks at selected loci. Top four tracks show ChIP-seq reads from the strain indicated as in Fig. 5. Genome map shows genes from 17978 (NZ_CP012004). Binding motif locations are denoted by black vertical lines (gray used in the case of one of two overlapping motifs). MACS2 peak intervals are shown as open rectangles.

Analysis of the ChIP-seq peaks from the *bfmRS* promoter and other representative sites validated by ChIP-qPCR revealed three general classes of BfmR binding activity. In one rare class, which included the promoter region upstream of *gidA*, significant FE was observed in the non-phosphorylatable *bfmR*(D58A) mutant along with the WT and Δ*bfmS* strains (Fig. 8A). Increased ChIP-seq reads were also seen at the *bfmRS* promoter in *bfmR*(D58A) mutant, although the FE was below the threshold of detection for peak calling (Fig. 8B). In a second category, which included the promoter regions upstream of *ompA*_Ab_ and ACX60_RS07610, binding was detected almost equivalently in both Δ*bfmS* and WT, but not in *bfmR*(D58A) (Fig. 8C). In a third category, exemplified by sites upstream of *pilM*, ACX60_RS01845 (Fig. 8D-E), *omp25* (Fig. 9A), and many others (eg, Fig. 10E), substantially higher BfmR binding was observed in the Δ*bfmS* strain background compared to that in the WT with no binding detected in *bfmR*(D58A). In several of the binding sites, clusters of multiple BfmR motifs were detected.

**Fig. 9.**
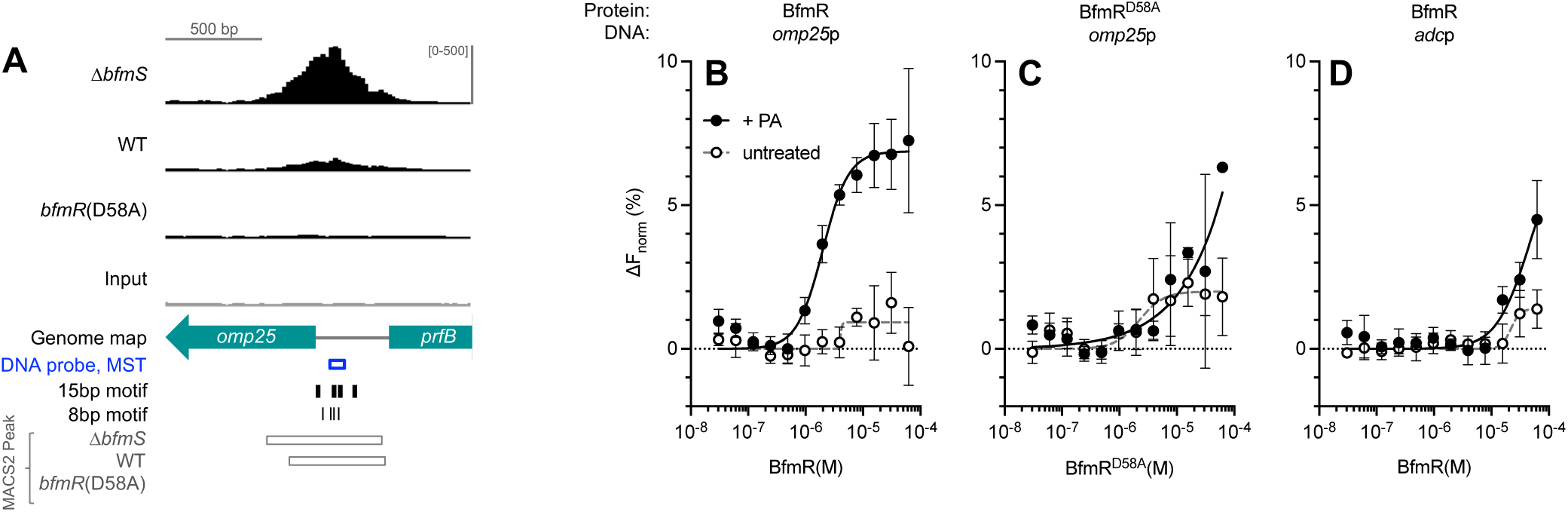
Phosphorylation enhances binding of BfmR to target DNA *in vitro*. (A) ChIP-seq reads aligning to the *omp25* promoter are depicted as in Fig. 8. (B-D) Results of MST assays. Binding of purified BfmR (B, D) or BfmR^D58A^ (C) to a fluorescein-labeled DNA probe was measured by MST. The DNA probe contained a 65 bp sequence from the promoter of *omp25* (B, C; indicated in blue in A) or *adc* (D). Data points show mean change in normalized fluorescence ± s.d. (n=3). Curves show nonlinear regression (specific binding with Hill Slope model).

**Fig. 10.**
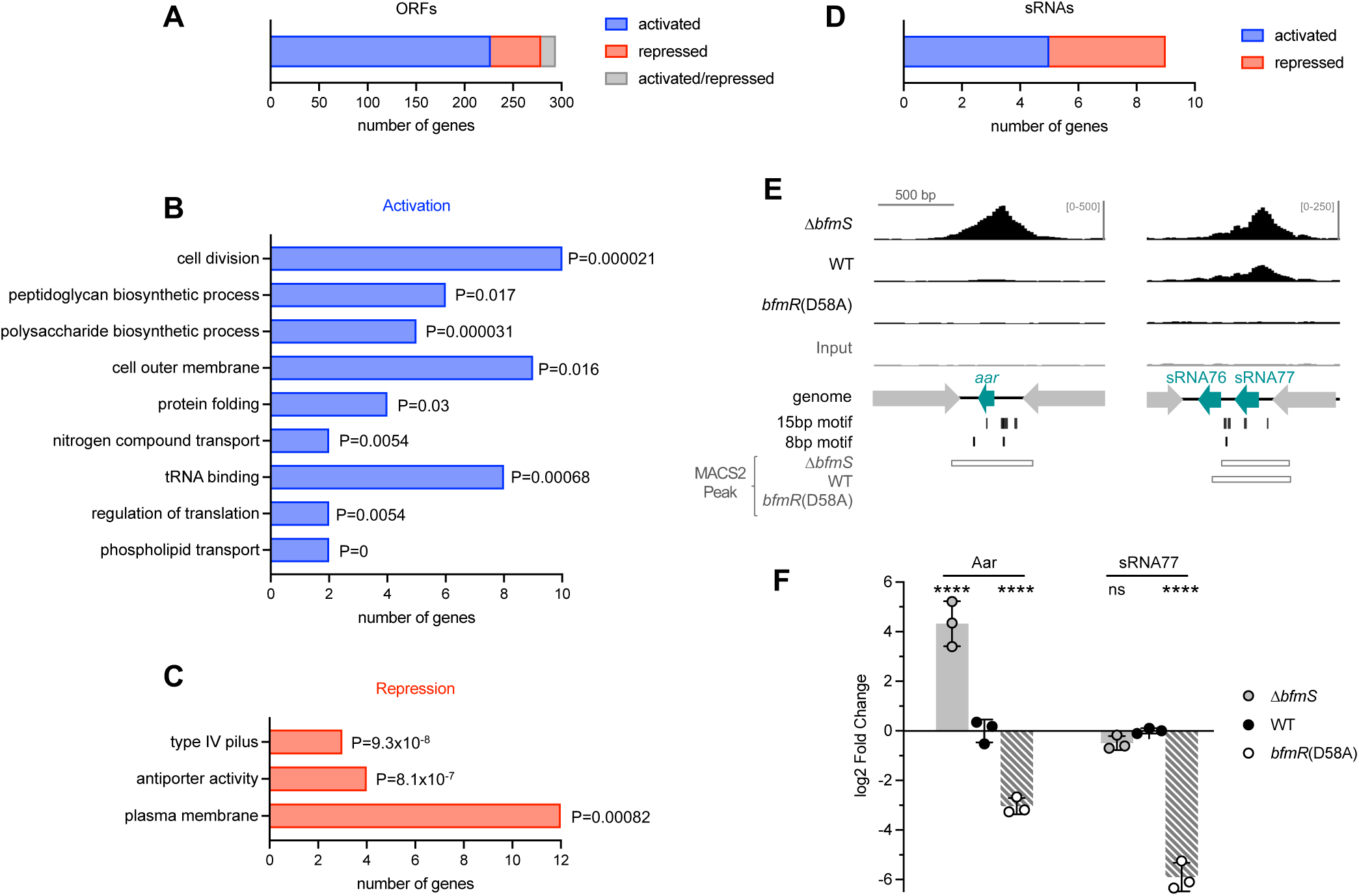
Analysis of the BfmR direct regulon. (A) Bar graph indicates the number of ORFs identified as direct targets of BfmR, with direction of regulation indicated by color. (B-C) GO term enrichment analysis of target ORFs directly activated (B) or repressed (C) by BfmR. A subset of the enriched GO terms and associated P-values (upper cumulative hypergeometric probability with Benjamini–Hochberg correction) are shown. (D) Bar graph shows the number of sRNAs identified as direct targets of BfmR activation or repression. (E) ChIP-seq reads aligning to the promoter regions of *aar* and *sRNA77* are depicted as in Fig. 8. (F) Gene expression in stationary phase cultures of the indicated strain was measured by RT-qPCR. Data show geomean of log2 fold change (vs WT) ± s.d. (n=3). Differences between mutant and WT were evaluated by two-way ANOVA with Dunnett’s multiple comparisons test. ns, P>0.05; ****, P≤0.0001.

To examine the phosphorylation-dependent binding of BfmR to newly identified target DNA containing the novel BfmR binding motif *in vitro*, we used MST (48). We again used the phosphodonor PA to generate phosphorylated BfmR (Fig. S2B) and combined the phosphorylated protein (or unphosphorylated control) with 5’ FAM-labeled DNA ligand. We first used a DNA sequence from the region upstream of *omp25*, just before its predicted −35 element, containing two 15bp BfmR binding motifs (each 15bp motif defined by one 8bp half-site) (Fig. 9A, Fig. S4A). When BfmR was phosphorylated, concentration-dependent binding to the *omp25*p DNA was observed, with an apparent Kd of 2 µM calculated from nonlinear regression analysis (Fig. 9B, +PA). By contrast, binding was not observed with unphosphorylated BfmR (Fig. 9B, untreated). Similar to the result with unphosphorylated WT BfmR, the BfmR^D58A^ mutant was defective for binding to the *omp25*p DNA despite the presence of PA treatment, showing only a partial change in MST signal at the highest protein concentrations (Fig. 9C). To assess the specificity of BfmR for DNA containing the identified binding motifs, an identically sized DNA ligand was designed with the sequence of a promoter, *adc*p, that lacked binding motifs and was not enriched in ChIP-seq. Compared to the affinity seen with *omp25*p, binding by phosphorylated BfmR to *adc*p was much weaker, with a curve fit yielding an apparent Kd of 45 µM (Fig. 9D). These results lend further support to the model that phosphorylation augments BfmR binding to target DNA.

### BfmR directly controls more than 300 genes including a set of sRNAs

To identify direct gene targets of BfmR from the ChIP-seq data, we integrated our results with information on genome-wide transcriptional regulation by BfmR. To this end, we interrogated published RNA-seq datasets examining the effects of deletions in *bfmRS* on gene expression in *A. baumannii* ATCC 17978 grown to exponential phase (6), as well as in ATCC 17961 grown to early stationary phase by Palethorpe et al (17). BfmR targets were identified using the following criteria: (1) the ORF closest to each BfmR ChIP-seq peak from NRA49 (Δ*bfmS*) was considered a target if it showed evidence of regulation, defined as a significant change (q < 0.05) in RNA-seq expression in any one of the *bfmRS* mutant strains compared to WT control. (2) For intergenic binding sites that occurred between a pair of divergent ORFs, the second-closest ORF was also considered a target if it was <400bp from the Peak summit and showed regulation by BfmRS as above. The list was manually curated to remove false positives, including instances in which the binding site was closer to a candidate sRNA than a coding sequence (see below). (3) Genes that form operons with the above loci were included as targets. This approach identified 294 ORFs as direct targets of BfmR (Fig. 10A, Dataset S3).

By analyzing the direction of the RNA-seq gene expression changes, genes were further classified as directly activated (233 ORFs) or repressed (54 ORFs) by BfmR (Fig. 10A). 15 genes showed evidence of both activation and repression, depending on the mutation analyzed by RNA-seq. For several genes, the location of the BfmR binding motifs relative to their predicted promoter was consistent with their classification as activated or repressed. For instance, in the *omp25*, *ACX60_RS01845*, and *slt* promoters, there are many BfmR binding sites upstream of the respective −35 elements, a location that could potentially enhance recruitment of RNA polymerase to the promoter (Fig. S4A), although this remains to be tested. With repressed genes *pilM* and *ACX60_RS07610*, a different arrangement is seen, with all the BfmR binding sites either overlapping or downstream of the predicted −10 element (Fig. S4B).

To assess the connection of the direct BfmR regulon to cellular functions, we identified its enriched GO terms (Materials and Methods). Among the genes directly activated by BfmR, GO terms related to cell division and biosynthesis of cell wall, capsule, and outer membrane were significantly enriched (adjusted p value <0.05) (Fig 10B, Dataset S4). This supports the notion that several of the up-regulated pathways implicated previously in envelope-linked phenotypes such as antibiotic resistance have elements that are under direct transcriptional control by BfmR (6). Interestingly, a prominent cellular component term that was enriched was the outer membrane, a reflection of the large number of outer membrane-localized proteins that are directly regulated by BfmR. Protein folding and translation terms were also enriched among the activated targets (Fig. 10B). In contrast to the activated pathways, targets that were directly repressed by BfmR were enriched for location in the plasma membrane, including integral membrane proteins that form a cation:proton antiport system (Pha/Mrp) as well as membrane proteins of the glutamate/aspartate transporter (GltJKL) and Type IV Pilus machinery (PilMNOP) (Fig. 10C, Dataset S4). BfmRS thus appears to reciprocally activate and repress production of proteins targeted for different cell envelope locales.

During examination of the location of ChIP-seq peaks, we noticed that some intergenic BfmR binding sites were unusual in that they resided between convergently transcribed genes (eg, peaks NRA49_chr-79, −145, and −222, Dataset S1, Fig. 10E). In the first of these examples, the binding site is in the intergenic DNA between the convergently oriented *trpS* and *sucD* ORFs (Fig. 10E). This is the exact locus that contains an sRNA gene, *aar*. Aar was first identified in *A. baylyi* as a putative regulator of amino acid metabolism (49). Its presence was recently confirmed in *A. baumannii*, in which it was shown to repress the *carO* outer membrane protein post-transcriptionally (50). This finding prompted an expanded search of the full set of *A. baumannii* sRNAs previously identified by high-resolution RNA-seq (29) for candidates directly regulated by BfmR. 11 sRNA loci, including *aar*, were identified that were in proximity (≤251bp) of a ChIP-seq binding site (Fig. S5). We then determined which sRNAs showed evidence of regulation by BfmR by reanalyzing the *bfmRS* mutant RNA-seq data to measure differential expression of sRNA genes (Materials and Methods). 9 of the 11 candidate sRNAs showed significant regulation, particularly in stationary phase, and these were deemed direct BfmR targets (Fig. 10D, Fig. S5, Dataset S3). 5 of these sRNAs showed activation by BfmR and 4 showed repression, based on pattern of expression across the *bfmRS* mutants compared to WT (Fig. 10D, Dataset S3). The annotation of three of these targets (Aar, SsrS, and sRNA77) as sRNAs had been validated previously by Northern blot (29,50). We confirmed direct regulation by BfmR for two of the validated sRNAs, Aar and sRNA77, by using two methods. First, BfmR binding to the promoters of *aar* and sRNA77 was verified by independent ChIP-qPCR, with BfmR phosphorylation associated with increased binding signal (see Fig. 6A). Second, to examine how phospho-BfmR affects sRNA transcription, we performed RT-qPCR on RNA extracted from *bfmR*(D58A), Δ*bfmS*, and control cultures in stationary phase, a condition known to increase Aar and sRNA77 levels (29,49,50). We found that the D58A mutation caused Aar and sRNA77 levels to decrease dramatically, by about 8-fold and 64-fold, respectively, compared to WT; while Δ*bfmS* caused Aar levels to increase 20-fold (Fig. 10F). These results are consistent with the model that BfmRS controls sRNA production as part of its broad regulatory program, particularly under stationary phase conditions.

## DISCUSSION

In this study we have demonstrated the critical role of BfmR phosphorylation in governing *A. baumannii* antibiotic resistance, virulence, and gene regulation at diverse loci. The non-phosphorylatable *bfmR*(D58A) mutant was defective in all these phenotypes. By analogy with other TCSs (51), BfmR is predicted to receive a phosphorylation signal in a regulated manner from its cognate receptor BfmS *in vivo*. BfmR phosphorylation appears to cause positive autoregulation, with *bfmR*(D58A) mutants showing lower BfmR protein levels compared to strains producing BfmR∼P. Despite correcting for lowered expression, BfmR^D58A^ had no ability to confer antibiotic resistance, yet it showed dominant-negative interference of BfmR^WT^ functions when highly overexpressed (Fig. 3). BfmR^D58A^ was also defective for phosphorylation, dimerization, and DNA-binding *in vitro*. These findings support Asp58 as the key site of phosphorylation and activation, in agreement with previous reports (17,23), and suggest that BfmR activates analogously to canonical OmpR/PhoB regulators (22). To uncover how phosphorylation affects BfmR binding to regulatory targets within bacteria, we performed ChIP-seq and integrated the resulting binding data with published RNA-seq gene expression profiles of BfmR. The results, revealing genome-wide dependence on BfmR∼P for DNA binding, agreed with the phenotypic and *in vitro* findings and facilitated definition of the comprehensive direct BfmR regulon. The regulon contains both previously identified targets as well as unappreciated factors, including novel BfmR-regulated sRNAs.

ChIP-seq yielded insights on where BfmR binds in the genome, its consensus target DNA motif, and the global role of phosphorylation in BfmR-DNA interactions. BfmR binding sites identified by ChIP-seq were commonly intergenic, consistent with direct regulation of promoter activity. A subset of sites were within coding regions, but these still tended to be close to ORF start codons. The role of intragenic binding remains unclear but could be compatible with regulation of transcription by DNA looping (52), roadblocking RNAP elongation (53), or modulating cryptic intragenic promoters (54). Heterogeneity was also observed across the ChIP FE values for binding sites in strains generating BfmR∼P. The wide dynamic range could reflect variable affinity for certain sequences or number of BfmR molecules bound at particular sites, although varied accessibility to immunoprecipitation due to presence of other proteins bound at certain sites is another possibility (55). Overall, higher levels of BfmR∼P (Δ*bfmS* > WT > *bfmR*(D58A)] yielded higher ChIP-seq FE, with almost all binding lost with BfmR^D58A^ (Fig. 5A-D), consistent with phosphorylation enhancing the interactions of BfmR with target DNA globally. Occupancy of the non-phosphorylated protein at a small number of promoters (such as *gidA* or *bfmRS*) is a possibility that requires additional verification. Future studies into how BfmR interacts with individual target sites will help evaluate the above models.

From the genome-wide ChIP-seq data, we determined the consensus DNA motifs targeted by BfmR. The most significant was a 15bp direct repeat sequence, and a second motif was an 8bp half-site comprising one of the repeats. For a DNA target containing the motifs (*omp25*p), we confirmed that phosphorylated, but not unphosphorylated, BfmR binds *in vitro* with low micromolar affinity (Fig. 9). The configuration of direct repeats with a 5bp spacer supports a model in which a BfmR dimer binds the recognition sequence with binding domains in a head-to-tail orientation, an arrangement seen with other response regulators (56,57). With both DNA motifs, a large proportion was located at the ChIP-seq peak center, consistent with mediating direct BfmR binding. Notably, there was a substantial proportion of binding sequences in which the 15bp motif was offset by about 30bp from peak center (Fig. 7C). The finding of this small offset could be explained by the possibility that this set of motifs is bound to larger multiprotein complexes involving BfmR, which would increase the length of DNA crosslinked and thus retrieved during ChIP (58). One possibility is that BfmR is contacting RNA polymerase at these sites, potentially reflecting RNAP recruitment and transcriptional activation of many targets (Fig. 10A) (52). BfmR could also be in a complex with other undetermined transcription factors. While an inverted arrangement of the same repeats identified in the *bfmRS* promoter (23) might play a role in BfmR autoregulation or other forms of control at some targets, the 15bp direct arrangement is widespread among diverse binding locations and is likely more relevant for genome-wide BfmR regulation.

The direct BfmR regulon was delineated in this work and comprises 303 genes (Fig. 10A,D). We have established that BfmR is global and bifunctional, activating many targets while acting as a repressor of others (Fig. 10). Among the activated proteins are K locus enzymes involved in capsule biosynthesis and protein glycosylation, and this direct control may contribute to the ability of *A. baumannii* to modulate virulence (10). Cell wall enzymes (eg, Slt, MurG, MurC, and Ddl), which are involved in intrinsic resistance to carbapenems (6,59), and a variety of outer membrane proteins with relevance to infection and drug resistance (eg, OmpA_Ab_, Omp25, CarO) (60–62), were also revealed as direct targets of activation. An additional cell envelope pathway directly activated by BfmR is periplasmic protein folding, which included two disulfide bond generation enzymes (DsbB and DsbC) (63), the chaperone SurA, and a DegP-family protease (ACX60_RS04935) (64). Upregulation of this pathway could contribute to how *A. baumannii* counteracts a major stress, defective periplasmic disulfide formation, shown previously to be a signal for BfmRS (9), and suggests the system may respond to other forms of protein misfolding in the envelope analogous to *E. coli* Cpx (65). In tandem, a separate group of cell envelope proteins was directly repressed by BfmR. These included inner membrane proteins mediating solute transport, as well as the PilMNOP complex involved in production of the type IV pilus. Downregulating these structures could represent part of the overall response to envelope protein folding stress (66). Repression of type IV pilus specifically could also allow *A. baumannii* to modulate motility, adherence, and/or horizontal gene transfer functions. Overall, direct reciprocal control of capsule (activated) and pili (repressed) would be expected to alter how the *A. baumannii* surface interacts with the environment, potentially contributing to the adaptability of the pathogen to different niches.

While the direct BfmR targets identified here help to disentangle its expansive regulon, hundreds of regulon members are assumed to be modulated by indirect mechanisms. Two examples include the β-lactamases *adc* and *oxa51*, whose transcription is dramatically altered in *bfmRS* mutants (6) but whose promoters were not detected bound to BfmR in ChIP-seq. For many indirect effects, it is possible that BfmRS acts through intermediate regulators to modulate gene expression. In this regard, we identified leads for two classes of intermediates. First, a number of transcriptional regulators were identified as direct targets of BfmR, including those known to alter the expression of large sets of genes from studies in *A. baumannii* (OmpR_Ab_, GacA (67,68)) or by analogy with other bacteria (IHF (69)). Second, we also identified a set of 9 sRNAs under the direct control of BfmR. The sRNAs include Aar, which was recently found to regulate CarO post-transcriptionally by blocking translation (50). The *carO* promoter is also directly targeted for transcriptional activation by BfmR (Dataset S3). Although the functions of most of the remaining sRNAs and several transcription factors remain to be defined, this suggests that BfmR uses multiple layers of control to tune expression of key targets.

In conclusion, these studies demonstrate the essential role of BfmR phosphorylation in *A. baumannii* virulence and antibiotic resistance and uncover mechanisms of global gene regulation controlled by this signal. Future work will examine how the phosphorylation signal is generated and regulated during homeostasis and stress, and how the activated protein functions to influence gene expression at distinct sites. In addition, work is needed to characterize the contribution of components of the direct BfmR regulon to modulating aspects of antibiotic resistance, host interactions, and growth in the context of infection. Interfering with the phosphorylation signal essential to the BfmR global regulatory network has the potential to contribute to strategies to potentiate drug activity and block the infectivity of the nosocomial pathogen.

## DATA AVAILABILITY

The raw sequencing data underlying this article are available in the NCBI Sequence Read Archive at https://www.ncbi.nlm.nih.gov/sra, and can be accessed with BioProject accession numbers PRJNA1129841 (BfmR ChIP-seq), PRJNA1129869 (ATCC 17978 *bfmRS* mutant RNA-seq), and PRJNA780533 (ATCC 17961 *bfmR* mutant RNA-seq). The differential gene expression data used in the article are available at https://doi.org/10.1371/journal.ppat.1007030 (ATCC 17978 *bfmRS* mutants), and https://doi.org/10.1128/jb.00494-21 (ATCC 17961 *bfmR* mutant). All other data are available in the article and in its online supplementary material.

## ACKNOWLEDGMENTS

We are grateful to Brandon Miller and Roman Manetsch for synthesis of phosphoramidate, Jinna Bai for constructing plasmids for *bfmR* overexpression, and members of the Geisinger laboratory for helpful discussions. We also thank Veronica Godoy and Kim Lewis for sharing equipment, and the Northeastern Institute for Chemical Imaging of Living Systems (CILS) core facility for providing access to MST and imaging instruments.

## FUNDING

This work was supported by Northeastern University startup funds; and the National Institutes of Health [grant number R01AI162996 to E.G.].

## Supplemental Files

**Fig. S1.** Supplemental information from Western blots.

**Fig. S2.** Analyses of BfmR *in vitro*.

**Fig. S3.** C-terminal 3XFLAG epitope tag preserves BfmR functions.

**Fig. S4.** Locations of BfmR binding motifs relative to promoters of example direct target genes.

**Fig. S5.** Analysis of sRNAs that are candidates for direct regulation by BfmR.

**Supplemental Table 1.** Bacterial strains, plasmids, and primers used in this study.

**Supplemental Data Set 1.** BfmR ChIP-seq peaks.

**Supplemental Data Set 2.** BfmR-binding motifs.

**Supplemental Data Set 3.** Direct BfmR targets.

**Supplemental Data Set 4.** GO Terms significantly enriched in the sets of genes activated or repressed by BfmR.

## Notes

### Competing Interest Statement

The authors have declared no competing interest.

### Summary of Updates

Revised the abstract and main text; updated supplemental material.

