## Supplemental Figures and Table for "A phosphorylation signal activates genome-wide transcriptional control by BfmR, the global regulator of *Acinetobacter* resistance and virulence"

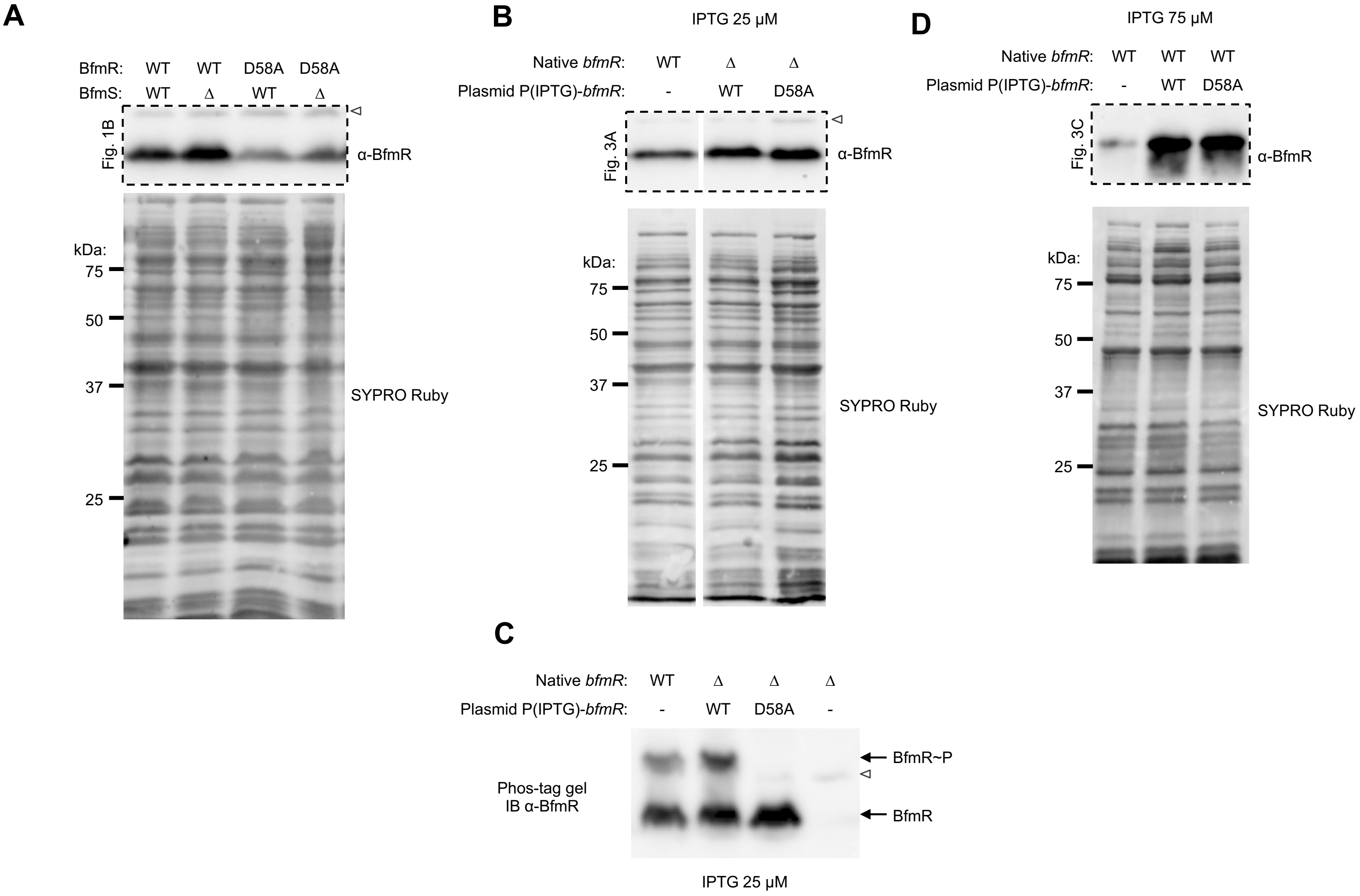

**Fig. S1. Supplemental information from Western blots.** (A, B and D) SYPRO staining of total protein in the blotted samples from Fig. 1B, 3A, and 3C, respectively. kDa values indicate the migration of molecular weight markers. (C) Phos-tag immunoblot of samples from Fig. 3A. Open arrowheads indicate non-specific protein band reacting weakly with BfmR antiserum.

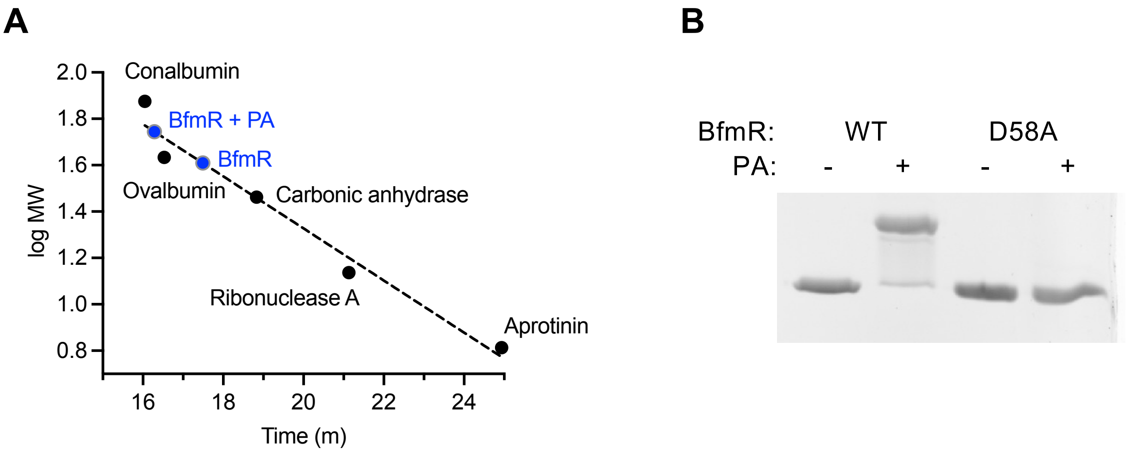

**Fig. S2. Analyses of BfmR *in vitro*.** (A) Estimation of BfmR molecular weight relative to protein standards without and with phosphorylation (Buffer S). Retention times of the protein standards (Gel Filtration LMW Calibration Kit, Cytiva) was plotted vs molecular weight. Estimated size of BfmR and BfmR+PA was determined by plotting their average peak retention time (n=3). (B) Efficient phosphorylation of purified BfmR protein used in MST experiments. BfmR was treated with or without PA in Buffer D, and phosphorylation analyzed by Phos-tag gel as in Fig. 4.

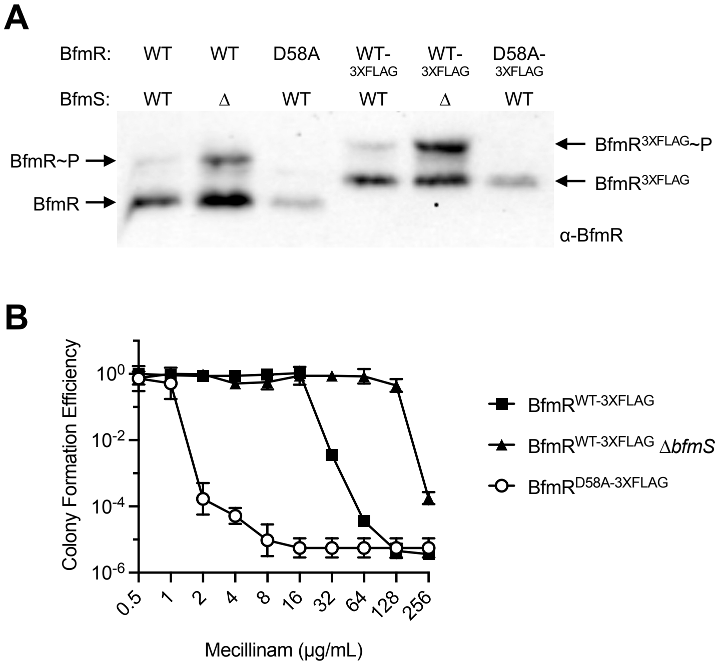

**Fig. S3. C-terminal 3XFLAG epitope tag preserves BfmR functions.** (A) Phos-tag Western blot analysis of BfmR phosphorylation in strains with the indicated *bfmRS* alleles. Blots were probed with BfmR antiserum. (B) Strains containing a C-terminal 3XFLAG tag on BfmR show levels of resistance to mecillinam resistance consistent with that seen with untagged strains [4]. Colony forming efficiency was measured on solid medium with mecillinam at the indicated dose compared to control medium lacking drug. Data points show geometric mean ± s.d. (n=3).

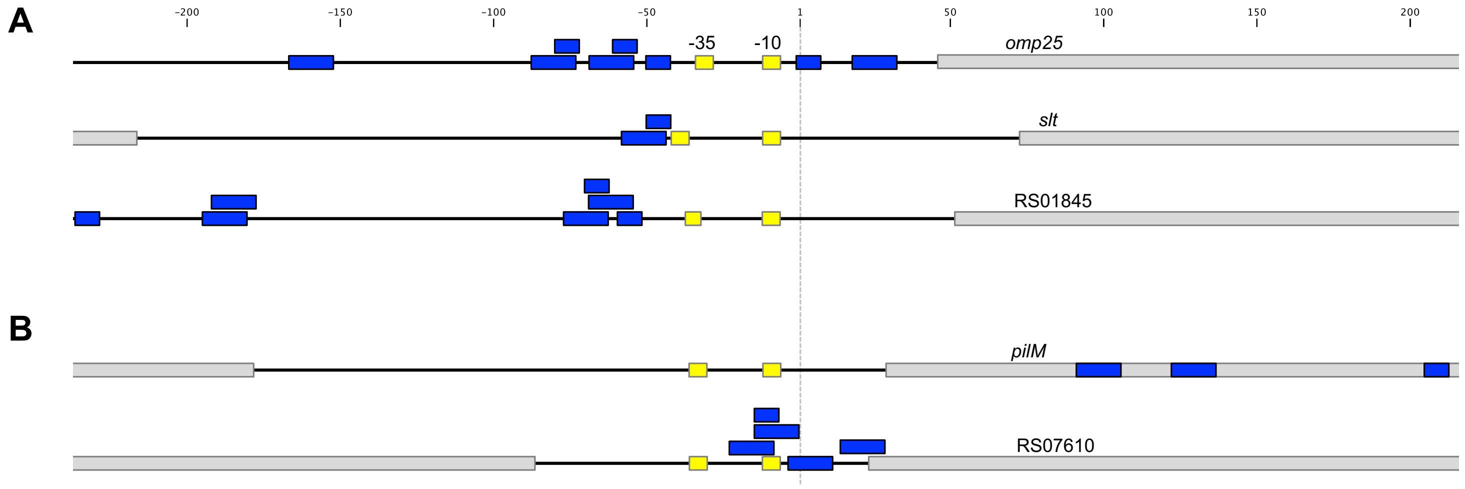

**Fig. S4. Locations of BfmR binding motifs relative to promoters of example direct target genes.** Promoter regions of activated (A) and repressed (B) targets are shown. Location of promoter elements was determined from [10] (*slt*, RS01845), [11] (*omp25*), or predicted by using BPROM software [12] (*pilM*, RS07610). -35 and -10 elements are depicted as yellow boxes. 15bp motif and 8bp BfmR binding motifs are shown as wide and narrow blue rectangles, respectively. ORFs are shown as grey rectangles. Dotted vertical line indicates the location of the TSS.

| Candidate target sRNA | | | ChIP-seq | | |
|---|---|---|---|---|---|
| ID | Name | Locus coordinates | Closest summit | Distance from sRNA | FE |
| sRNA21 | Aar | 833848 - 833746 | 833899 | -51 | 10.39 |
| sRNA84 | 6S/SsrS | 2772408 - 2772589 | 2772420 | 12 | 3.91 |
| sRNA77 | | 2544991 - 2544831 | 2544991 | 0 | 7.11 |
| sRNA103 | | 3560294 - 3560386 | 3560284 | -10 | 5.91 |
| sRNA40 | | 1714402 - 1714497 | 1714456 | 54 | 3.8 |
| sRNA29 | | 1381844 - 1381668 | 1382030 | -186 | 2.77 |
| sRNA54 | | 2180417 - 2180273 | 2180601 | -184 | 2.86 |
| sRNA76 | | 2544740 - 2544594 | 2544991 | -251 | 7.11 |
| sRNA87 | | 2841274 - 2841173 | 2841524 | -250 | 2.25 |
| sRNA30 | | 1396374 - 1396483 | 1396224 | -150 | 5.14 |
| sRNA85 | | 2774620 - 2774564 | 2774684 | -64 | 4.09 |

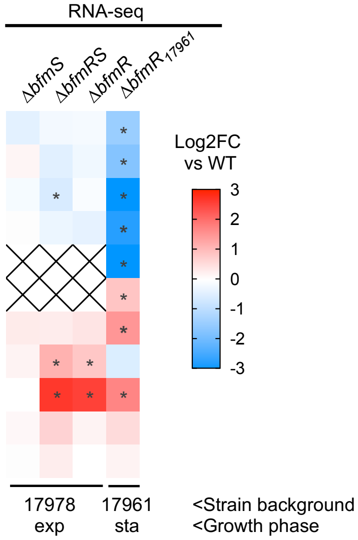

**Fig S5: Analysis of sRNAs that are candidates for direct regulation by BfmR.** sRNA loci were identified that are in proximity to a BfmR binding site identified by ChIP-seq. Locus coordinates indicate the position of the sRNA gene in 17978 (NZ_CP012004). FE indicates ChIP-seq fold enrichment in NRA49 (Δ*bfmS*). Heatmap shows log2 fold change in RNA-seq reads, comparing the indicated mutant to WT. X indicates raw read count was too low to analyze. *, adjusted p-value <0.05; exp, exponential phase; sta, onset of stationary phase. Gray shading indicates candidates not showing significant BfmRS-dependent change in gene expression in RNAseq.

### Table S1. Strains, plasmids, and primers used in this study

#### Strains

| Designation | Genotype or description | Strain ID | Reference |
|---|---|---|---|
| ***A. baumannii*** | | | |
| 17978 | cerebrospinal fluid isolate, AbaAL44$^+$ ("UN") type, ATCC | EGA83 | [1, 2] |
| 17978 Δ*bfmS* | ATCC 17978 Δ*bfmS*::*aacC1* | EGA195 | [3] |
| 17978 Δ*bfmRS* | ATCC 17978 Δ*bfmRS*::*aacC1* | EGA495 | [4] |
| 17978 Δ*bfmR* | 17978 Δ*bfmR*::*aacC1* | EGA496 | [4] |
| 17978 *bfmR*(D58A) | 17978 *bfmR*(D58A) | NRA407 | This work |
| 17978 *bfmR*(D58A) Δ*bfmS* | 17978 *bfmR*(D58A) Δ*bfmS* | NRA446 | This work |
| 17978 *slt*p-GFP | EGA83 with pEGE315 | NRA480 | This work |
| 17978 *bfmR*(D58A) *slt*p-GFP | NRA407 with pEGE315 | NRA481 | This work |
| 17978 Δ*bfmRS slt*p-GFP | EGA495 with pEGE315 | NRA482 | This work |
| 17978 *pilM*p-GFP | EGA83 with pNRE216 | NRA433 | This work |
| 17978 Δ*bfmS pilM*p-GFP | EGA195 with pNRE216 | NRA435 | This work |
| 17978 *bfmR*(D58A) *pilM*p-GFP | NRA407 with pNRE216 | NRA437 | This work |
| 17978 Δ*bfmR* P(IPTG)-*bfmR* | EGA496 with pJE86 | NRA460 | This work |
| 17978 Δ*bfmR* P(IPTG)-*bfmR*(D58A) | EGA496 with pNRE138 | NRA461 | This work |
| 17978 Δ*bfmR* vector | EGA496 with pEGE305 | NRA462 | This work |
| 17978 P(IPTG)-*bfmR* | EGA83 with pJE86 | NRA147 | This work |
| 17978 P(IPTG)-*bfmR*(D58A) | EGA83 with pNRE138 | NRA456 | This work |
| 17978 vector | EGA83 with pEGE305 | NRA365 | This work |
| 17978 *bfmR*-3xFLAG | 17978 *bfmR*-3xFLAG | NRA28 | This work |
| 17978 *bfmR*(D58A)-3xFLAG | 17978 *bfmR*(D58A)-3xFLAG | NRA29 | This work |
| 17978 *bfmR*-3xFLAG ΔbfmS | 17978 *bfmR*-3xFLAG Δ*bfmS* | NRA49 | This work |
| 17978 Δ*gtr6* | 17978 Δ*gtr6* | JBA202 | [5] |
| 17978 Δ*gtr6 bfmR*(D58A) | 17978 Δ*gtr6* bfmR(D58A) | NRA486 | This work |
| ***E. coli*** | | | |
| DH5α | *supE44* Δ*lacU169* (ϕ80lacZΔM15) *hsdR17 recA1 endA1 gyrA96 thi*-1 *relA1* | EGE1 | [6] |
| DH5λpir | DH5α (λpir) *tet*::Mu *recA* | EGE4 | [7] |
| BL21 (DE3) pLysS | F- *ompT hsdSB*(rB- mB-) *gal dcm* (DE3) pLysS (Cm$^r$) | NRE81 | Novagen |

#### Plasmids

| Plasmid | Description | Reference |
|---|---|---|
| pUC18 | *oriColE1* MCS Cb$^r$ | [8] |
| pEGE305 | P(IPTG) shuttle vector (*ori-pBR322 ori*-pWH1277 *bla*::*lacI*$^q$-T5*lac*P Tc$^r$) | [4] |
| pJB4648 | Conditionally replicating allele exchange plasmid (*oriTRP4 oriR6K sacB* Gm$^r$) | [9] |
| pNRE157 | pUC18 containing homology arms for introducing *bfmR*(D58A) allele | This work |
| pNRE169 | pJB4648 containing homology arms for introducing *bfmR*(D58A) allele | This work |
| pEGE245 | reporter plasmid with promoterless *gfp*mut3 (*ori*-pBR322 *ori*-pWH1277, Tc$^r$) | [4] |
| pEGE315 | *slt*p-GFP reporter plasmid (*ori*-pBR322 *ori*-pWH1277, Tc$^r$) | [4] |
| pNRE211 | pUC18::*pilM*p | This work |
| pNRE216 | pEGE245::*pilM*p-GFP | This work |

**Table S1 (continued)**

| | | |
|---|---|---|
| pJE83 | pUC18::*bfmR* | This work |
| pJE86 | pEGE305::*bfmR* | This work |
| pNRE137 | pUC18::*bfmR*(D58A) | This work |
| pNRE138 | pEGE305::*bfmR*(D58A) | This work |
| pNRE80 | pUC18::[NdeI]*bfmR*[BamHI] | This work |
| pNRE85 | pET28a::*bfmR* | This work |
| pNRE127 | pET28a::*bfmR*(D58A) | This work |
| pEGE228 | pUC18 containing *bfmR*-3xFLAG-*bfmS* allelic exchange construct | This work |
| pNRE26 | pJB4648 containing *bfmR*-3xFLAG-*bfmS* allelic exchange construct | This work |
| pNRE12 | pUC18 containing *bfmR*(D58A)-3xFLAG-bfmS allelic exchange construct | This work |
| pNRE27 | pJB4648 containing *bfmR*(D58A)-3xFLAG-bfmS allelic exchange construct | This work |
| pNRE31 | pUC18 containing upstream homology arm for *bfmS* deletion with *bfmR*-3xFLAG | This work |
| pNRE32 | pUC18 containing upstream homology arm for *bfmS* deletion with *bfmR*(D58A)-3xFLAG | This work |
| pNRE33 | pUC18 containing downstream homology arm for *bfmS* deletion | This work |
| pNRE34 | pJB4648 containing homology arms for Δ*bfmS* deletion with bfmR-3xFLAG | This work |

**Oligonucleotide primers**

| **Primer name** | **Sequence (5' – 3'; restriction site underlined if present)** | **RE site(s)** |
|---|---|---|
| ***Allelic exchange*** | | |
| bfmR-D58x-R | AGACCACAAGATCCGGTTGCTC | |
| bfmR-D58A-F | TGGCTGTCATGTTGCCGGGTGC | |
| dbfmS-down-F | CATGTCGACGCAATTGCCCATGATGAACT | SalI |
| dbfmS-down-R | CATGGTACCTTTAAACAACCGCCATTAAAGACC | KpnI |
| dbfmS-up-F | TATGGTACCACTGTGTTTAAACACTCGACCAACC | KpnI |
| dbfmS-up-R | ATCGGATCCAGTTTGGTGAACGCCTACTTGT | BamHI |
| BamHI-500up-BfmRD58A-F | ACATGGATCCCGGTAGATCAATCTTGACTTT | BamHI |
| SalI-500dwn-BfmRD58A-R | ATGTGTCGACGATTTTACAATCCATTGGTTTCTTTAAC | SalI |
| ***Reporter/bfmR expression*** | | |
| SacI-pilMp-Fwd | ATGTGAGCTCGATCAAAAAAATTGGACGCACG | SacI |
| KpnI-pilMp-Rev | TAGCGGTACCACTATTGTCCTATTATTTTTTATCCCC | KpnI |
| bfmRS-ecoF | GTGGAATTCGCAAATGATAAACGAATGTATCTGCAAG | EcoRI |
| bfmRonly-pstR | TTACTGCAGCGACCAACCTTATAGGAAGTTTAATCAG | PstI |
| NdeI-BfmR-Fwd | ACTGCCATATGAGCCAAGAAGAAAAG | NdeI |
| BamHI-BfmR-Rev | CAGTGGATCCTTACAATCCATTGGTTTCTTTAAC | BamHI |
| ***ChIP-qPCR*** | | |
| dnaA-qF1 | GTAGATTCTCGTCCTGGTAGTATTT | |
| dnaA-qR1 | CCTTAGCAGGTTGAGGTATAGG | |
| bfmR qPCR FWD Set 2 | TCGTCGTCTTCAACGATCAGAA | |
| bfmR qPCR REV Set 2 | GCAAATGATAAACGAATGTATCTGCAAG | |
| surA Set 1F | CTATGCCTATGCGCCATACAA | |
| surA Set 1R | CATGACCATAGAAGCGGTAAGG | |
| 01845 Set 2F | GTTGTTCATGTGATACATGCCTAT | |

**Table S1 (continued)**

| | |
|---|---|
| 01845 Set 2R | AGCAATTATTATGCCGTTTCCTC |
| itrA Set 1F | GACCGTACCAGAAACAGCAT |
| itrA Set 1R | TCGCCGCAAAGGTTTACA |
| 18040 Set 1F | CATCTTCGCGCTGCCTATAA |
| 18040 Set 1R | ATTGCAGAATTTGTACCGCTATTAC |
| wzi Set 1F | ACAGCCGATGAAGCAGTT |
| wzi Set 1R | ACTTTGAGAATTGTGCTGACAT |
| omp25 Set 1F | AGCACATGGTTACAGTCCAG |
| omp25 Set 1R | TGTTACAACTTTCAGTCTAGAGCA |
| ompA Set 2F | TCAAGCACTTGGAAAGTCTATCA |
| ompA Set 2R | TTGTTGTTCAAGCTCAGCCTA |
| pilM/pbp1a Set 1F | GGACAATAGTGTGCTCAGGTTAT |
| pilM/pbp1a Set 1R | CTTGACAGAGAGCTCTAACAACTTA |
| gidA Set 1F | TGGCTTACCGTAAATCATGTCA |
| gidA Set 1R | CGCCACCGATAACGATAACA |
| putP Set 2F | GCGACCTGGATTAGGTTACAA |
| putP Set 2R | TGTAGCACGGTAGGCAAATAA |
| efp Set 2F | AACCGGGTAAAGGCCAAG |
| efp Set 2R | CGCCATCGTTGTATAGGTAGTT |
| aar-ChIP qPCR-1F | GGTGATCACTGCGTAGAACAA |
| aar-ChIP qPCR-1R | GCGTCACTAATATAACTTGAGTAGGT |
| RS07610-ChIP qPCR-2F | CGCTTGGCTAATGTTGTTAGTC |
| RS07610-ChIP qPCR-2R | GCTCATTATCTAAATCGACACTTACTC |
| sRNA77-ChIP qPCR-4F | GTGGATCGAGGAGATATTACGATTAC |
| sRNA77-ChIP qPCR-4R | ACCCAAATGGCGTCGAAA |
| ***RT-qPCR*** | |
| rpoC-qF4 | CAAACGGTGAGCCAATCATC |
| rpoC-qR4 | GCCTTCACCTTTCGCATTT |
| pilM-qPCR 2F | GCTCTCTGTCAAGAACGGTAAA |
| pilM-qPCR 2R | CTGCAACTGCTTCTGGATTTAAG |
| slt70-qF1 | CACTAGGCCGTTTAGCAAATAAT |
| slt70-qR1 | GGCTACGGTTCGATAGAGATAC |
| aar qPCR FWD Set 1 | TGATATGAACCTCACGACATTTCT |
| aar qPCR REV Set 1 | GGTGATCACTGCGTAGAACAA |
| sRNA77-qPCR-1F | ATTTGCTCTTTGCTAGCTGTTT |
| sRNA77-qPCR-1R | GGTAATCGTAATATCTCCTCGATCC |
| ***MST*** | |
| omp25-FB-FAM | /56-FAM/ATATATTAAATTAAATATAGTTACATAAAAAGCACATGGTTACAGTCCAGTTACTTGGACAAGAT |
| omp25-RB | ATCTTGTCCAAGTAACTGGACTGTAACCATGTGCTTTTTATGTAACTATATTTAATTTAATATAT |
| adc-FAM-F | /56-FAM/TATTTAAAAAGAAAGATGCCTACTTTTATAACAAAAATCACCTAATTTAATTGTTATGTTTTATA |
| adc-R | TATAAAACATAACAATTAAATTAGGTGATTTTTGTTATAAAAGTAGGCATCTTTCTTTTTAAATA |
